# Structure–Property–Performance Engineering of Hydrogel Depots for Long-Acting Peptide Delivery

**DOI:** 10.64898/2026.05.17.725768

**Authors:** Changxin Dong, Andrea I. d’Aquino, Samya Sen, Alakesh Alakesh, Carolyn K. Jons, Noah Eckman, Christian M. Williams, Leslee T. Nguyen, Jerry Yan, Olivia M. Saouaf, Ye Eun Song, Ian A. Hall, Katie Lu, Manoj K. Manna, Sara Kapasi, Sai Anahitha Kottamasu, Tatum Wilhelm, Vannessa Doulames, John H. Klich, Wencke Reineking, Eric A. Appel

**Affiliations:** Department of Materials Science & Engineering, Stanford University, Stanford, CA 94305, USA; Department of Mechanical Engineering, Indian Institute of Technology Madras, Chennai, Tamil Nadu, 600036, India; Department of Chemical Engineering, Stanford University, Stanford, CA 94305, USA; Department of Biochemistry, Stanford University, Palo Alto, CA 94305, USA; Department of Bioengineering, Stanford University, Stanford, CA 94305, USA; Department of Biology, Stanford University, Stanford, CA 94305, USA; Wallace H. Coulter Department of Biomedical Engineering, Georgia Institute of Technology, Atlanta, GA 30332, USA; Veterinary Service Center, Department of Comparative Medicine, Stanford University School of Medicine, Stanford, CA 94305, USA; ChEM-H Institute, Stanford University, Stanford, CA 94305, USA; Department of Pediatrics (Endocrinology), Stanford University, Stanford, CA 94305, USA; Woods Institute for the Environment, Stanford University, Stanford, CA 94305, USA

**Keywords:** GLP-1, hydrogel, sustained delivery, diabetes, weight management

## Abstract

Controlled release systems for subcutaneous peptide delivery often exhibit a pronounced initial burst release followed by inadequate maintenance of therapeutic exposure, limiting depot lifetime and increasing pharmacokinetic variability. Here, we engineer a dynamic, injectable hydrogel depot technology for months-long delivery of lipidated peptides. Using semaglutide as a model, we establish a modular formulation framework integrating: (i) formulation-driven tuning of depot mechanics to control release kinetics, (ii) cargo complexation strategies leveraging hydrophobic and multivalent ion-mediated interactions, and (iii) oxidative stabilization through sacrificial antioxidant excipients. We evaluated depot performance by rheology, in vitro cargo release, and in vivo pharmacokinetic and pharmacodynamic studies in rodents. Optimized formulations sustained semaglutide exposure for over six weeks from a single administration with two-fold reduction in peak-to-trough exposure and comparable total bioavailability relative to daily dosing, resulting in improved glucose control, weight regulation, and preservation of pancreatic islet content. These results suggest potential for quarterly dosing in humans. Together, this work establishes integrated and generalizable structure-property-performance relationships that account for cargo–matrix and cargo–excipient interactions across burst, diffusion, and erosion regimes to inform a practical formulation framework for engineering long-acting depots for sustained peptide delivery.

## 1. Introduction

Peptide therapeutics represent a rapidly expanding class of biopharmaceuticals for the treatment of metabolic disease, cancer, and other chronic conditions.^1,2^ Yet, many of these drug molecules exhibit short systemic half-lives that necessitate frequent dosing to maintain therapeutic exposure, leading to poor patient adherence and significant peak-to-trough fluctuations that can drive adverse side effects.^1,3^ Several newer peptide therapeutics, including glucagon-like peptide-1 receptor agonists (GLP-1 RAs) such as Semaglutide (Sema; Ozempic®, Wegovy®, and Rybelsus®; Novo Nordisk) or Tirzepatide (Mounjaro® and Zepbound®; Lilly), bear lipid modifications that imbue these molecules with longer half-lives that support weekly dosing;^4–6^ however, weekly dosing is still too burdensome for patients to maintain compliance.^7,8^

One solution to the burden of frequent dosing is the development of sustained delivery depot technologies.^9–14^ The earliest generation of these long-acting delivery technologies was based on biodegradable polymer microparticles and enabled sustained release for select peptide drugs, exemplified by leuprolide drug product formulations such as Lupron Depot®.^15,16^ Unfortunately, these systems have exhibited success with only a small set of peptide therapeutics on account of manufacturing and delivery challenges including low drug loading efficiency that restricts accessible doses and wastes precious drug, poor stability during encapsulation which often requires exposure to organic solvents, and poor control over the duration of drug release.^17,18^ As a result, many peptide therapeutics cannot be effectively formulated using these sorts of solid polymer matrices.^19^ To address this technological gap, injectable hydrogel depot formulations have emerged as versatile platforms for long-acting drug formulations, enabling sustained release through dynamic network rearrangement and slow dissolution of the hydrogel.^9–11^ Yet, prior work on the development of hydrogel depot technologies, including our own, has been limited by challenging manufacturing, poor peptide bioavailability, and/or high peak-to-trough exposure variability that can often drive undesirable side effects.^11,20–24^ Moreover, truly long-term peptide release over the timeframe of months with injectable hydrogel depots remains challenging due to burst release, a mis-match in cargo delivery and depot erosion, and chemical instability of the peptides during prolonged residence *in vivo*.^13,25–28^ Addressing these limitations requires coordinated control over hydrogel mechanics, cargo–matrix interactions, and peptide stability.

In this work, we introduce a distinct injectable hydrogel depot technology that overcomes these limitations and use this platform to develop a design framework for engineering interaction-based hydrogel depots with extended-release lifetimes and improved pharmacokinetic profiles. We have previously reported the development of a hydrogel platform using hydrophobically modified cellulose polymers bearing pendant stearyl chains (HPMC-C_18_) that form transient physical crosslinks via micelle-mediated association of hydrophobic side chains.^29^ These transient micellar junctions impart shear-thinning and self-healing behavior that enables minimally invasive injection followed by reformation of a stable subcutaneous depot.^30^ We build on this work to develop three complementary formulation design strategies to engineer long-acting peptide depots: (i) hydrogel vehicle formulation to tune micellar crosslink density and bulk mechanical properties, (ii) cargo complexation to modulate intermolecular interactions and effective diffusion within the matrix, and (iii) oxidation-prevention strategies such as sacrificial antioxidant excipients to protect sensitive residues and stabilize the peptide during residence in the depot. Each strategy was evaluated for hydrogel rheological properties, *in vitro* release kinetics, and pharmacokinetics in murine models.

We apply these design approaches using Sema as a model peptide therapeutic. The saturated C_18_ diacid fatty acid moiety on the peptide^31^ promotes selective association with the hydrophobic micellar domains within our HPMC-C_18_ hydrogel system, enabling intrinsic cargo-matrix coupling that retards diffusion and prolongs depot residence time. As a key example of the increasingly important class of peptide therapeutics,^32,33^ Sema enables systematic interrogation of hydrophobic interaction-driven depot formation and release control to identify broadly applicable design principles for lipidated and amphiphilic peptide therapeutics. We conceptualize release from these systems as transport through a dynamically evolving network in which cargo association and network relaxation co-determine flux. This framework yields three distinct transport regimes, including (i) burst, (ii) diffusion, and (iii) erosion, whose relative contributions can be tuned through orthogonal formulation parameters. We demonstrate how this approach enables prolonged systemic exposure with reduced peak-to-trough variability and extended depot persistence, establishing a modular strategy for engineering long-acting peptide depots for chronic disease treatment.

## 2. Results and Discussion

### 2.1. Impacts of formulation on dynamic hydrogel crosslinking

Hydroxypropyl methylcellulose (HPMC) derivatives bearing stearyl (C_18_) side chains are commercially established polymers for which hydrophobic modification is simple, scalable, and reproducible, with broad use in hydrophobic drug solubilization and topical formulations.^34,35^ We have previously demonstrated that the grafted C_18_ moieties associate into transient nanodomains that act as physical crosslinks between the HPMC polymer chains in aqueous media.^29,30,36^ In these assemblies, C_18_-rich micellar cores are surrounded by solvated HPMC backbones, forming dynamic core-corona junctions leading to viscoelastic network formation.^37^ These reversible, non-covalent junctions drive formation of injectable, self-healing hydrogels without chemical crosslinkers. Hydrophobic moieties such as the lipid modifications on many commercial peptides readily partition into the C_18_-associated domains, modulating junction stability while preserving dynamic exchange.^29,38^ As a result, HPMC-C_18_ hydrogel formulations can be generated that exhibit pronounced shear thinning and rapid self-healing, allowing facile injectability followed by reformation of a localized subcutaneous depot capable of sustained drug release ideal for chronic disease management (**Fig. 1a**).

**Fig. 1.**
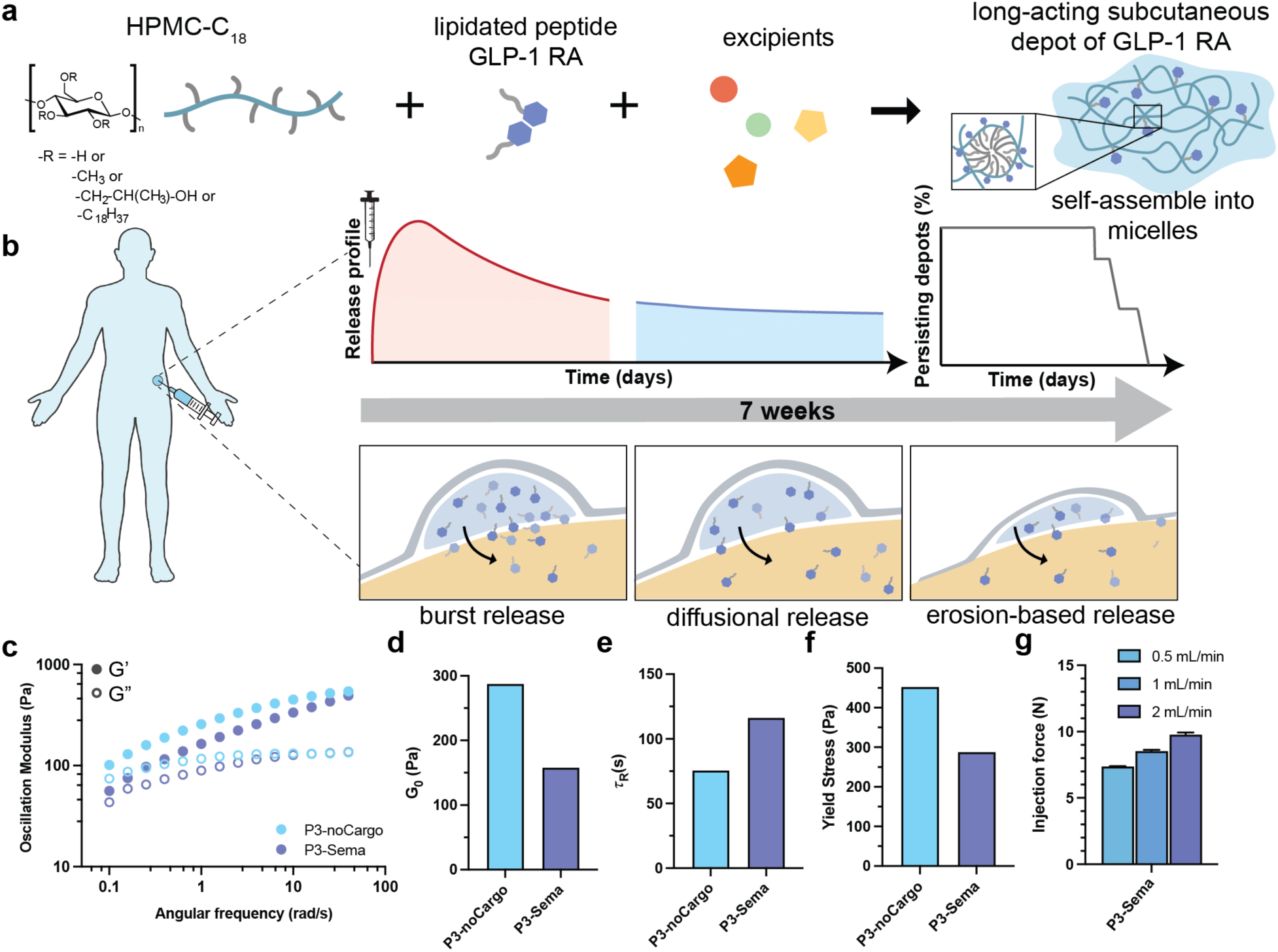
Summary of hydrogel-based subcutaneous depot design and release behavior. **(a)** Lipidated GLP-1 receptor agonists associate with hydrophobically modified cellulosics like HPMC-C_18_ to form a self-assembled subcutaneous depot, with functional excipients incorporated to modulate long-acting release. **(b)** Conceptual schematic illustrating the three distinct release regimes governing drug transport from injectable hydrogel depots: burst, diffusion-dominated, and erosion-mediated. The release profile exhibits three distinct regimes: (i) initial burst release, (ii) diffusion-governed release, and (iii) a network erosion-dominated release. **(c)** Oscillatory frequency sweep of 3 wt% HPMC-C_18_ with 1 mg/mL semaglutide (denoted P3-Sema) and without cargo (denoted P3-noCargo). **(d)** Plateau modulus (G₀) comparison between P3-noCargo and P3-Sema. **(e)** Relaxation time (τ_R_) extracted from oscillatory shear rheological measurements. **(f)** Dynamic yield stress obtained from Herschel-Bulkley fits to steady shear rheological data (**Fig. S2a**). **(g)** Injection force for P3-Sema at flow rates of 0.5, 1, and 2 mL/min through a 40 mm-long 21-gauge needle.

Drug release from HPMC-C_18_ depots proceeds through three characteristic regimes (**Fig. 1b**): (i) an initial burst regime reflecting rapid release of loosely associated cargo, (ii) an intermediate diffusion-dominated release regime that sustains systemic exposure, and (iii) a prolonged erosion-mediated regime as the depot dissolves at the injection site. To modulate these release behaviors, 3 wt% HPMC-C_18_ (denoted P3) was selected as the baseline hydrogel formulation, promoting formation of a more persistent and mechanically robust depot while enabling systematic tuning of excipient–cargo interactions. This composition was chosen to mitigate the initial burst and prolong the diffusion-dominated release phase. All subsequent studies therefore employ 3 wt% HPMC-C_18_ as the reference formulation.

Rheological characterization demonstrated that formulations with and without Sema (denoted P3-Sema and P3-noCargo, respectively) form solid-like networks with weak frequency dependence across the probed range (**Fig. 1c–f**), consistent with physically crosslinked hydrogels suitable for subcutaneous depots. Incorporation of semaglutide modestly decreases the oscillatory moduli (**Fig. 1c**), resulting in a lower plateau modulus G′₀ (**Fig. 1d**, **Supplementary Discussion 1**) and decreased dynamic yield stress (**Fig. 1f**), but similar flow properties (**Supplementary Figure 1**).^39–41^ These changes suggest partial softening of the hydrophobic junctions upon incorporation of the lipidated peptides, consistent with Sema partitioning into micellar domains, leading to a marginally softer network that would nevertheless be expected to exhibit similar injectability. Notably, the relaxation time τR increases in the presence of Sema (**Fig. 1e**), indicating slower junctional exchange and more persistent network dynamics. Thus, peptide loading yields a slightly softer yet mechanically coherent gel that retains solid-like character and well-defined yielding behavior desirable for depot formation and sustained cargo delivery.

Owing to this modest softening, P3-Sema remains readily injectable, despite the slower crosslink exchange dynamics. Injection forces measured through a 21-gauge needle at flow rates between 0.5 and 2 mL/min fall within a 50 N range, which is a practical range for subcutaneous administration (**Fig. 1g**, **Supplementary Figure 2**).^38,42^ The Sema/HPMC-C_18_ association supports formation of a self-assembled depot that maintains sufficient mechanical integrity to localize at the injection site while enabling long-acting release.

#### 2.1.1 Influence of surfactant-like excipients

Sema (Ozempic®, Wegovy®, and Rybelsus®) was selected as the therapeutic cargo for sustained delivery from our hydrogel depot materials owing to its fatty acid modification and extended plasma half-life relative to other GLP-1 RAs.^43–45^ Because drug release from HPMC-C_18_ hydrogels is governed not only by bulk stiffness but also by micellar crosslink dynamics, understanding how surfactant-mediated perturbation of hydrophobic junctions alters network mechanics and release behavior provides mechanistic would provide insight to inform rational formulation design and delivery strategy optimization. We have previously shown that nonionic surfactants, including Tween 20 (Tw20) or Tween 80 (Tw80), insert into the hydrophobic junctions, thereby accelerating crosslink exchange without altering the crosslink density and therefore the plateau modulus, resulting in reduced burst release for non-lipidated macromolecular cargos such as human IgG (hIgG).^36^ In HPMC-C_18_ depots loaded with Sema (**Fig. 2a**), the effect of Tw80 differed markedly from prior observations with hIgG,^36^ potentially reflecting differences in hydrodynamic dimensions as Sema oligomers exhibit an average hydrodynamic diameter (d_H_) of ∼ 4.6 nm, which is substantially smaller than hIgG (d_H_ ∼ 10-12 nm). To further probe this size-dependent behavior, dynamic light scattering measurements reveal that addition of Tw80 increased the apparent hydrodynamic diameter of Sema assemblies relative to Sema alone, consistent with surfactant interaction with C_18_-rich hydrophobic domains (**Fig. 2b**).

**Fig. 2.**
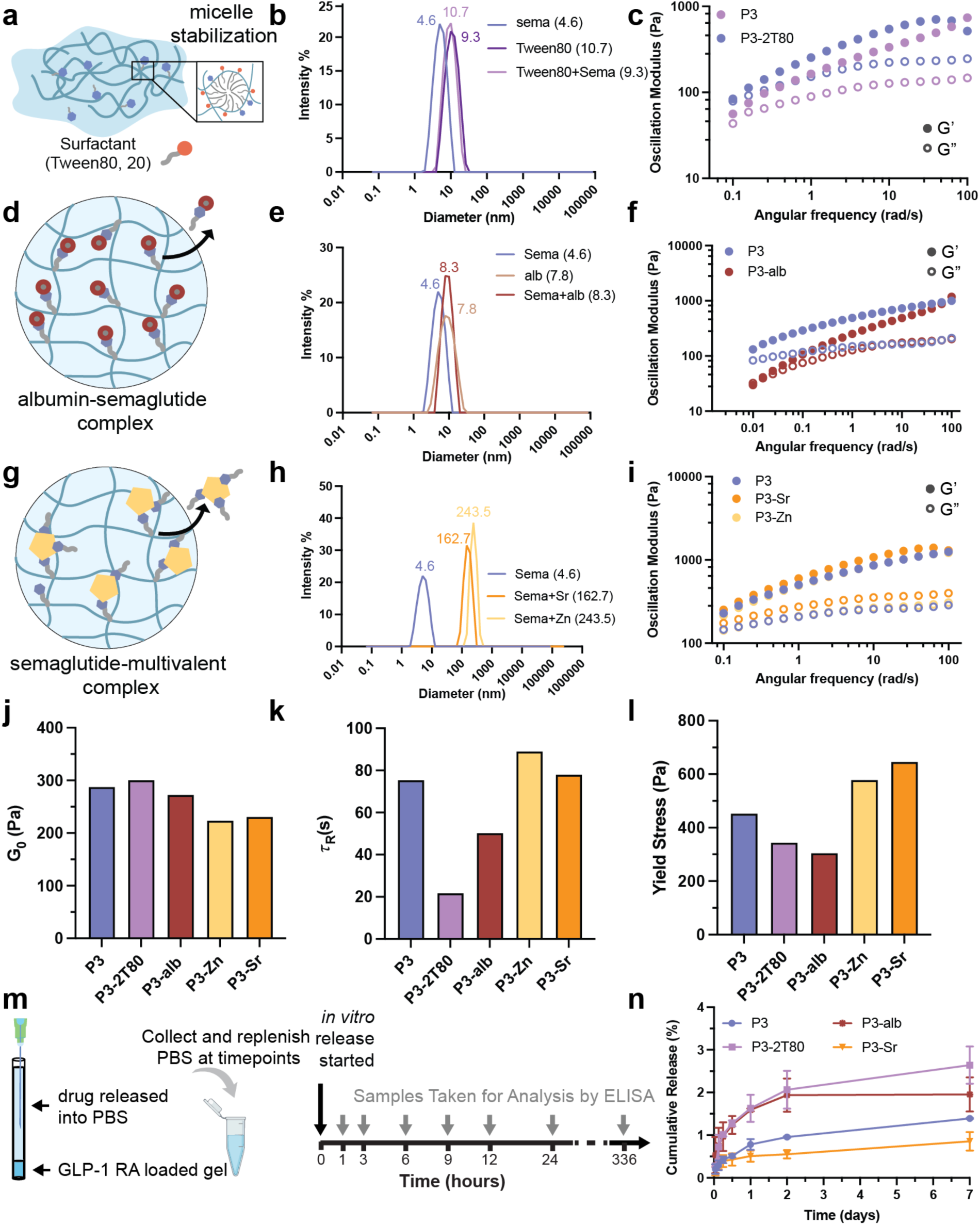
Cargo size modification tunes excipient-semaglutide interactions and governs hydrogel crosslinking mechanics and cargo release. **(a)** Schematic illustrating surfactant-mediated micelle stabilization of semaglutide in the presence of Tween-20. **(b)** Dynamic light scattering (DLS) size distributions of Sema with and without Tween-20, indicating formation of larger peptide-surfactant assemblies. **(c)** Oscillatory frequency sweeps of 3 wt% HPMC-C_18_ (P3) hydrogels with Sema or Tween-20-Sema, revealing excipient-dependent changes in viscoelastic response. **(d)** Schematic of albumin-semaglutide complex formation. **(e)** DLS size distributions of semaglutide incubated with albumin, demonstrating formation of higher-molecular-weight complexes. **(f)** Frequency sweeps of P3 and albumin-loaded P3 hydrogels (P3-alb). **(g)** Schematic illustrating divalent cation (Sr²⁺)-mediated multimerization of semaglutide. **(h)** DLS size distributions showing Sr²⁺-induced peptide multimer formation. **(i)** Frequency sweeps of Sr²⁺-containing P3 hydrogels highlighting altered network mechanics. **(j–l)** Summary of plateau modulus (G₀), relaxation time (τ_R_), and dynamic shear yield stress for formulations including P3, P3-alb, and P3-Sr (**Supplementary Discussion 1**). **(m)** Schematic of *in vitro* release setup; PBS is replenished at each time point and released GLP-1 RA is quantified by ELISA (**Supplementary Discussion 2**). **(n)** Representative cumulative release profiles over 7 days for P3 and P3-T80 formulations in diffusion-dominated release behavior.

Incorporation of Sema into the P3-Sema hydrogels also resulted in a modest reduction in storage modulus relative to the cargo-free P3-noCargo hydrogels (**Fig. 1c**), indicating partial plasticization of the micellar crosslinking junctions. Notably, this softening effect was comparable to that observed in Tw80-containing formulations (**Fig. 2c**), suggesting that Sema itself acts as a weak amphiphile capable of associating with the C_18_ micellar crosslinks without substantially destabilizing the network. The lipidated tail of Sema promotes insertion into the hydrophobic micellar cores, while the peptide backbone remains solvated in the surrounding aqueous phase. This dual affinity enables Sema to participate in micellar clustering without disrupting the junctional structure. As a result, Sema reinforces hydrophobic association within the network rather than weakening it, rendering additional surfactant unnecessary, affirming the suitability of Sema as model therapeutic cargo for evaluation of the outlined formulation strategies for sustained delivery.

Comparison of formulations with and without Tw80 (P3-T80 and P3, respectively) indicates that HPMC-C_18_ is sufficient to sustain long-acting delivery of lipidated peptide cargos without auxiliary surfactants (**Fig. 3c** and **3f-i**). Inclusion of Tw80 reduced network stiffness and accelerated systemic decline in serum concentrations, leading to lower overall exposure. These results suggest that surfactant-mediated perturbation of the associative network compromises depot performance, highlighting that excipients must be evaluated not only for their impact on the therapeutic cargo but also for their impact on network integrity. The C_18_ lipid modification of Sema promotes co-assembly within the hydrophobic micellar junctions of the HPMC-C_18_ matrix. The peptide therefore contributes to network organization rather than behaving as a freely diffusing solute. Competitive disruption of micellar clustering by surfactant addition weakens these hydrophobic associations and diminishes pharmacokinetic control. This behavior contrasts with that of hydrophilic protein cargos such as IgG, which do not partition into hydrophobic micellar domains and may require surfactants to modulate network relaxation.^36^ These findings support a mechanistic framework in which fatty-acid-modified peptides are inherently compatible with C_18_-based supramolecular depots, enabling formulation simplicity without the need for surfactant additives.

**Fig. 3.**
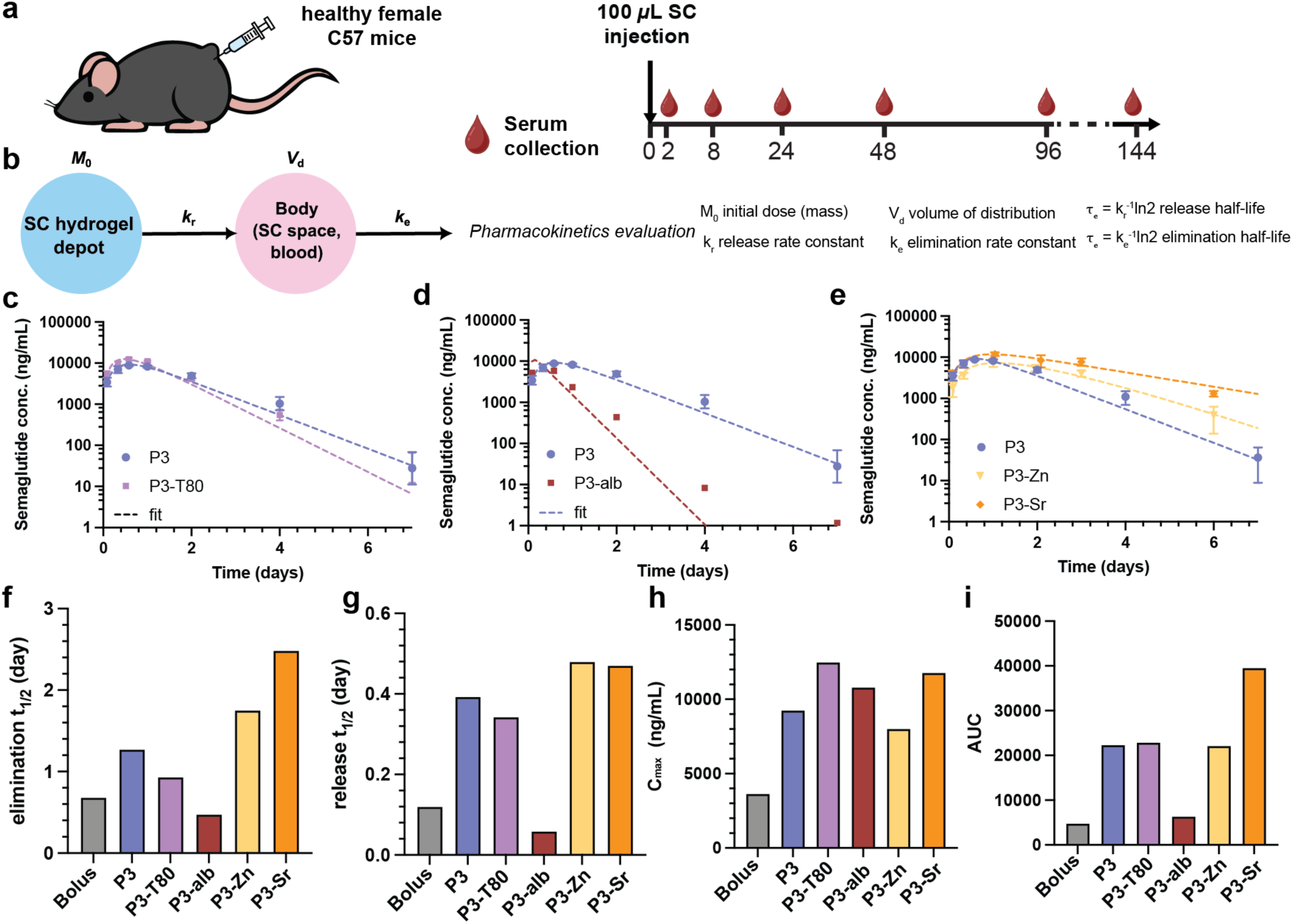
Pharmacokinetic evaluation of semaglutide hydrogel depots in healthy C57BL/6 mice. **(a)** Schematic of the pharmacokinetics study design in mice. Eight-week-old female C57BL/6 mice received a single 100 µL subcutaneous hydrogel injection. Serum was collected at 2, 8, 12, 24, 48, and 144 hours for pharmacokinetic profiling. Data were fit to a two-compartment model incorporating the SC depot and systemic compartment to extract release and elimination parameters (**Supplementary Discussion 3**). **(b–d)** Semaglutide serum concentration–time curves for P3 and modified P3 formulations, including Tween-80 (P3-T80), albumin prebinding (P3-alb), and Zn²⁺-complexed formulations (P3-Zn). **(e)** Elimination half-life (t_1/2_, elimination), **(f)** release half-life (t_1/2_, release), **(g)** C_max_, and **(h)** AUC for all conditions. For **(f–i)**, parameters were obtained from a combined fit across the full dataset. These metrics highlight how excipient-dependent semaglutide clustering modulates burst release and sustained exposure *in vivo*.

#### 2.1.2. Prebinding of peptides to albumin increases effective cargo size

Sema’s C_18_ modification enables high-affinity albumin binding (**Fig. 2d**), making pre-complexation a viable strategy for increasing cargo size to enhance entrapment in the hydrogel matrix for slower diffusion and release. Although DLS confirmed modest enlargement of Sema multimers upon albumin binding (d_H_ ∼ 8.3 nm versus 4.6 nm; albumin alone 7.8 nm; **Fig. 2e**), albumin incorporation destabilized the hydrogel network. Indeed, gel stiffness decreased (*i.e.*, reduced plateau modulus) and early-phase release accelerated, indicating disruption of hydrophobic micellar junctions. A possible explanation is competition of albumin with C_18_ domains for micellization through fatty-acid binding pockets and hydrophobic surface patches, perturbing associative crosslinks between the cellulosic chains. This destabilization was more pronounced than with Tw80, consistent with the observed increase in burst release and reduced pharmacokinetic performance (**Fig. 2e–h, Supplementary Figure 2b**). Beyond network effects, pre-complexation with albumin adds manufacturing complexity and regulatory burden. Despite increasing apparent cargo size, albumin pre-binding does not provide a practical advantage for depot design in this system. More broadly, these findings underscore that excipient selection must be evaluated not only for its impact on cargo size but also for its influence on network rheology. Indeed, even modest molecular interactions can substantially alter micellar crosslinking, stress relaxation, and ultimately negatively impact release behavior. Mechanistically, albumin likely accelerates release through two coupled effects: (i) sequestration of a fraction of Sema into a more mobile albumin-bound pool, reducing its participation in hydrophobic junctions, and (ii) direct reduction of micellar crosslink density, thereby lowering the stiffness of the hydrogels and facilitating faster stress relaxation. These observations indicate that increasing cargo size alone is insufficient to improve cargo retention in the depot as there may be an associated increase in network relaxation dynamics and therefore faster diffusive and erosive release.

#### 2.1.3. Multivalent ion complexation increases cargo size

Sema contains multiple anionic residues capable of coordinating divalent cations. We hypothesized that addition of Zn^2+^ or Sr^2+^ would induce peptide clustering (**Fig. 2g**), producing aggregates substantially larger than free peptide (**Fig. 2h**). Zn^2+^ generated the largest complexes (d_H_ ∼ 244 nm), consistent with stronger coordination to histidine and carboxylate groups, whereas Sr^2+^ formed smaller clusters (d_H_ ∼ 163 nm). Analysis of plateau modulus (G₀), relaxation time (τ_R_), and yield stress further indicated enhanced resistance to flow and prolonged stress relaxation for Sr^2+^ and Zn^2+^ formulations, which indicates a more persistent depot and slower diffusive release (**Fig. 2j–l**). The increased frequency-dependent moduli for both ions indicate increased network stiffness, which is desirable for prolonged cargo residence within the depots (**Fig. 2i–l, Supplementary Figure 2c**). *In vitro* data measuring the rate of Sema release from these gel formulations reflected size-dependent effects. Increasing effective cargo dimension reduced early burst and lowered cumulative release over 7 days, with the strongest attenuation in burst release observed for the multivalent systems (**Fig. 2n**). This suggests that cargo hydrodynamic size serves as an independent and tunable parameter governing peptide transport within the supramolecular hydrogel depots.

### 2.2. Pharmacokinetic analysis in mice

To characterize Sema pharmacokinetics in the various hydrogel formulations described above and to avoid variability from uneven type 2 diabetes induction, we used healthy 8-week-old female C57BL/6 mice and administered a single subcutaneous hydrogel dose. Serum samples were collected at 2, 8, 12, 24, 48, and 144 hours post-injection (**Fig. 3**, **Supplementary Discussion 2**). Pharmacokinetic profiles were fit to a two-compartment model (SC depot and systemic compartment). Elimination half-life (elimination t_1/2_), release half-life (release t_1/2_), C_max_, t_max_, and AUC were used to characterize *in vivo* release (**Supplementary Discussion 3**). Elimination half-life reflects the time required for serum Sema concentration to drop by half and release half-life reflects the time required for half of the depot-bound semaglutide to be released. C_max_ and t_max_ represent the peak Sema serum concentration and time to peak, and AUC captures total systemic exposure. These parameters extracted from the pharmacokinetics profile inform formulations with the lowest burst release and most sustained delivery.

While Zn^2+^ produced large aggregates, tighter coordination may hinder dissociation and limit systemic availability, as evident from short half-lives of both release and elimination. In contrast, the Sr^2+^ formulation (P3-Sr) exhibited prolonged elimination half-life, comparable release half-life, and nearly two-fold higher area under the curve (AUC) relative to Zn^2+^ depots (**Fig. 3e–h**). These findings suggest that moderately labile, charge-mediated clustering can yield superior systemic exposure compared to stronger metal-peptide coordination. Cargo size augmentation is effective only when balanced with reversible dissociation kinetics, highlighting coordination strength as a critical design variable for sustained-release peptide depots.

Commercial parenteral Sema formulations (Ozempic^®^ and Wegovy^®^) are formulated at pH ∼ 7.4. Lowering the pH induces Sema clustering by making semaglutide more hydrophobic, which increases the cargo size and effectively lowers C_max_ in the mice pharmacokinetics profile. The pharmacokinetic profiles of P3 formulated in different buffer systems reveal clear pH-dependent release characteristics (**Supplementary Tables 1–2**). With a phosphate buffer at pH 7.4, P3 formulations exhibited a shorter release half-life (t_1/2_ = 0.40 days) and faster absorption, reaching t_max_ at 0.61 days and a C_max_ of 9240 ng/mL. However, the total drug exposure (AUC = 22,300 ng day mL⁻¹) was lower, indicating a more rapid initial release followed by faster clearance (elimination t_1/2_ = 1.27 days). With a citrate buffer at pH 4, P3 formulations demonstrated a longer release half-life (t_1/2_ = 1.05 days) and delayed t_max_ (1.07 days), accompanied by a comparable C_max_ (8740 ng/mL) but a higher AUC (29,600 ng day mL⁻¹). Yet, elimination half-life remained similar (t_1/2_ = 1.17 days), suggesting that the extended exposure was primarily due to slower release rather than altered systemic clearance.

The difference in release behavior can be attributed to the effect of pH on Sema’s ionization and hydrophobicity, and therefore its association behavior. The isoelectric point (pl) of Sema is approximately 5.4,^46^ so at pH < 5.4 the peptide carries a more positive net charge and exhibits increased hydrophobicity due to protonation of ionizable groups, promoting stronger association with the hydrophobic domains of the HPMC-C_18_ network. This enhanced hydrophobic interaction improves peptide retention within the gel matrix, resulting in slower diffusion, reduced burst release, and a more sustained release profile. In contrast, at pH > 5.4 Sema carries a net negative charge; therefore, under physiological conditions (pH 7.4) the peptide is negatively charged. Acidic buffering (pH < 5.4) thus improves formulation stability and release control from a pharmacokinetic standpoint. Additional studies are needed to determine whether therapeutic efficacy can be fully retained in the treatment of T2D. Nevertheless, pH modulation represents a practical strategy to fine-tune peptide hydrophobicity and intermolecular interactions, providing an additional lever to modulate pharmacokinetic behavior.

### 2.3. Combating oxidation effects in subcutaneous depots

Sema is susceptible to oxidative degradation in storage, *in vivo* in a depot, during serum sample storage, and analytical quantification. Oxidation-prone residues including His7, Tyr10, and Trp31 are particularly vulnerable to modification that can disrupt structural integrity and receptor-binding activity.^47,48^ Tryptophan (Trp31) is highly susceptible to oxidative cleavage or indole-ring modification, which can alter protein stability and fluorescence properties. Histidine (His7) is reported to readily react with reactive oxygen species (ROS) to form oxo-histidine or imidazoline derivatives, potentially distorting binding-site geometry and impairing receptor activation.^49^ Tyrosine (Tyr10) may undergo oxidation to form tyrosyl radicals via phenol oxidation, which influence receptor binding affinity and peptide activity. To mitigate these degradative pathways, we introduced sacrificial antioxidants including L-methionine and α-tocopherol into the hydrogel formulations (**Fig. 4a**).^49,50^ These antioxidants are expected to preferentially react with ROS within the subcutaneous depot, thereby reducing oxidative stress on the encapsulated peptide during sustained release. Importantly, incorporation of methionine did not measurably alter the rheological properties of the hydrogel network, as confirmed by dynamic flow sweep measurements (**Supplementary Figures 2d and 3**), indicating that any observed differences in release behavior would arise from antioxidant–peptide interactions rather than changes in bulk material mechanics.

**Fig. 4.**
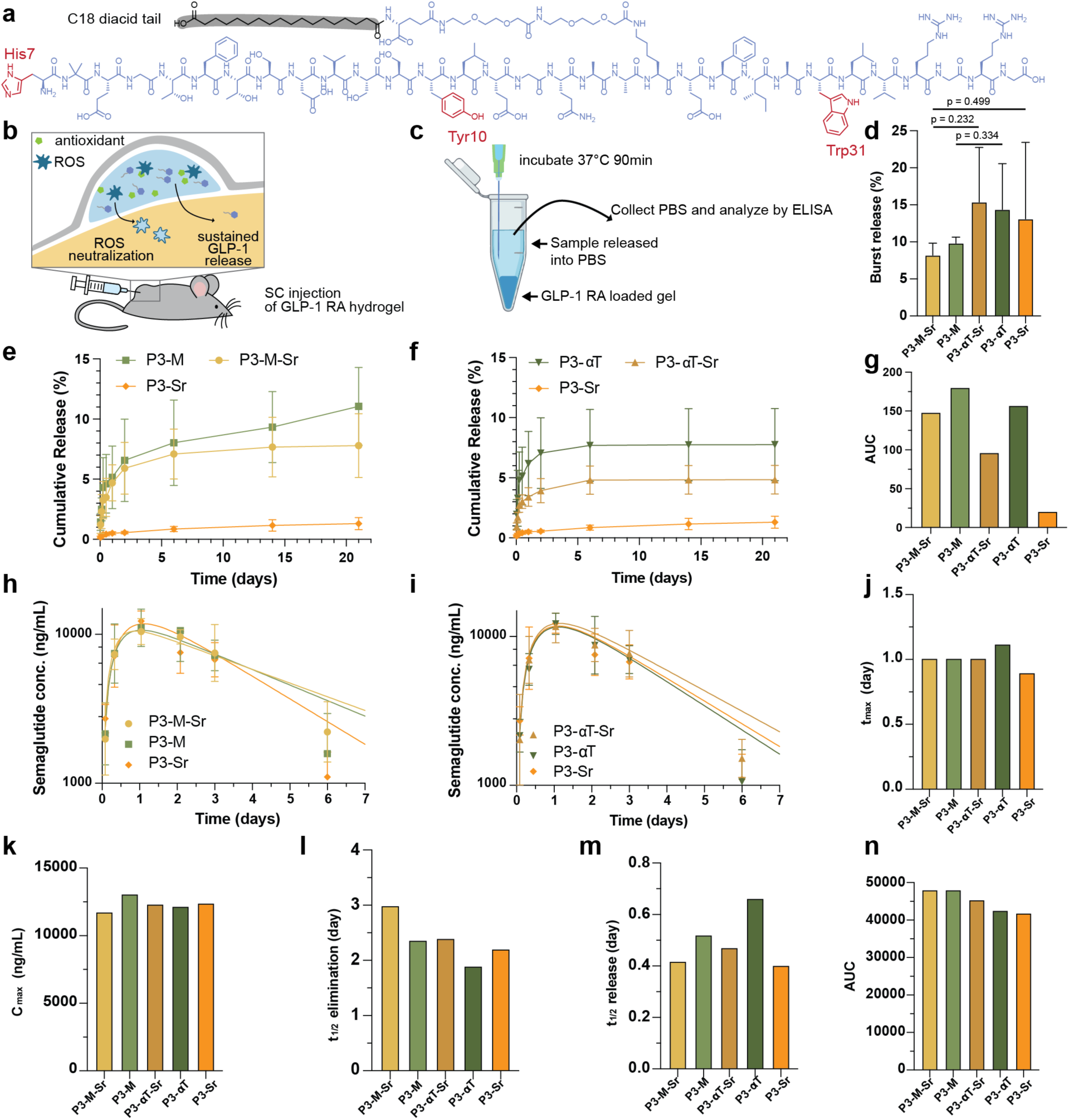
Strategies to mitigate semaglutide oxidation in hydrogel depots. **(a)** Semaglutide structure highlighting oxidation-prone residues (His7, Tyr10, Trp31) and schematic of antioxidant incorporation into the hydrogel depot. L-methionine (M, hydrophilic sacrificial antioxidant) and α-tocopherol (αT, hydrophobic antioxidant) were evaluated. **(b)** Capillary burst-release assay used to quantify early-stage release of semaglutide. **(c)** Burst-release comparison across formulations showing reduced initial release with methionine-containing depots. **(d-f)** Short-timescale release kinetics for semaglutide and antioxidant-modified formulations. Statistical comparisons in **(d)** were performed using a two-tailed Welch’s t-test (n = 3). **(g–h)** Serum pharmacokinetics in healthy mice for P3-M-Sr and P3-αT-Sr, fit with a two-compartment model (**Supplementary Discussion 3**). **(i–m)** Extracted PK parameters, including elimination half-life, release half-life, lag time, C_max_, and AUC. Antioxidant excipients reduce oxidative degradation, suppress burst release, and enhance sustained semaglutide exposure *in vivo*. Model parameters **(g), (j), (k), (l), (m)** and **(n)** were obtained from a combined fit across the full dataset for each formulation.

L-methionine (M) and α-tocopherol (αT) were selected as model antioxidants representing distinct radical-scavenging mechanisms. α-Tocopherol is a lipid-soluble chain-breaking antioxidant exhibiting high rate constants for peroxyl radical scavenging (10⁶–10⁷ M⁻¹ s⁻¹), consistent with its established role in suppressing lipid peroxidation.^50^ In contrast, methionine primarily undergoes sacrificial oxidation to sulfoxide and does not function as an efficient chain-terminating lipid radical scavenger.^46^ Their oxidation pathways reflect their relative reactivity, where methionine is converted to its sulfoxide and sulfone forms, while α-tocopherol undergoes multi-step oxidative degradation. Importantly, methionine is hydrophilic whereas α-tocopherol is hydrophobic, which affects how each excipient partitions within the hierarchical structure of the depot. Because the P3-Sr formulation showed the most favorable pharmacokinetics with the highest t_max_ of 0.997 days, lowest C_max_ of 11800 ng/mL, and highest AUC of 39500 ng day mL⁻¹ (**Fig. 3e–i**), P3-Sr was selected as the base formulation to further optimize formulation based on antioxidant-API interactions. In **Fig. 4**, combined strategies of antioxidant and complexation were further compared in both *in vitro* and *in vivo* experiments. To resolve early burst-release behavior that is not readily distinguishable in long-term capillary experiments, we established a short-timescale burst-release assay (**Fig. 4c, d**). While capillary release measurements capture long-term mass transport under surface-area-limited and quiescent conditions, they lack temporal resolution during the initial diffusion-dominated phase, where loosely associated cargo is rapidly liberated and released. In the burst release assay, Sema-loaded hydrogels were incubated at 37 °C for 90 min, and released peptide was quantified by ELISA. This complementary experimental approach enables quantitative assessment of formulation-dependent differences during the early release window that may be obscured in extended capillary measurements.

Within the first 90 min of incubation, methionine-containing formulations (P3-M-Sr and P3-M) exhibited reduced peptide release relative to P3-Sr, P3-αT, and P3-αT-Sr. Although mean release values were lower in the methionine-containing groups, statistical significance was not reached under nonparametric analysis (p > 0.05), likely due to variability within small cohorts (n = 3). In contrast, 21-day capillary release profiles revealed greater cumulative exposure, reflected by increased area under the curve (AUC), for antioxidant-containing groups, including P3-M and P3-αT, consistent with sustained peptide availability over extended durations (**Fig. 4e–g**). The combination of attenuated early release and enhanced cumulative release supports a dual function of methionine in suppressing burst release while promoting prolonged delivery. These *in vitro* data indicate that incorporating antioxidants, particularly methionine, improves release control and sustained exposure. The discrepancy between methionine and αT may additionally arise from differences in solvent optimization and excipient solubilization.

The four antioxidant-containing hydrogel formulations were subsequently evaluated in healthy female C57BL/6 mice using the same pharmacokinetic analysis protocol described above (**Fig. 4a, b**), with P3-Sr serving as the reference control (**Fig. 4g, h**). P3-αT exhibited the longest time to maximum concentration (t_max_ = 0.997 days; **Fig. 4j**). P3-M-Sr achieved the lowest peak concentration (C_max_ = 10,500 ng/mL; **Fig. 4k**) and the longest elimination half-life (t_1/2_ = 2.34 days; **Fig. 4m**). Notably, P3-M-Sr and P3-M yielded the highest overall exposure, with AUC values of 39,540 and 40,500 ng day mL⁻¹, respectively. Based on the observed favorable balance between reduced peak exposure (C_max_) and enhanced cumulative exposure (AUC), P3-M-Sr and P3-M were advanced to the next stage for further evaluation, in the T2D rat model, underscoring the functional contribution of methionine and the added benefit of Sr^2+^ incorporation.

### 2.4. Pharmacokinetic analysis in rats with type 2 diabetes

We evaluated how methionine-containing hydrogels control long-term Sema exposure in a T2D rat model (**Fig. 5**). The 42-day study design (**Fig. 5a**) allows us to follow both early and late phases of release as the injected depot volume in rats is sufficiently large to ensure persistence for the full study duration. This experimental design allows us to clearly resolve both diffusion- and erosion-driven elimination components using a two-mode, two-compartment PK modeling strategy (**Fig. 5b, Discussion S4**) that enables calculation of additional PK parameters, including the peak-to-trough ratio, which reflects therapeutic stability by comparing C_max_ to the steady-state plateau. A lower peak-to-trough ratio corresponds to reduced burst release and more sustained peptide delivery.

**Fig. 5.**
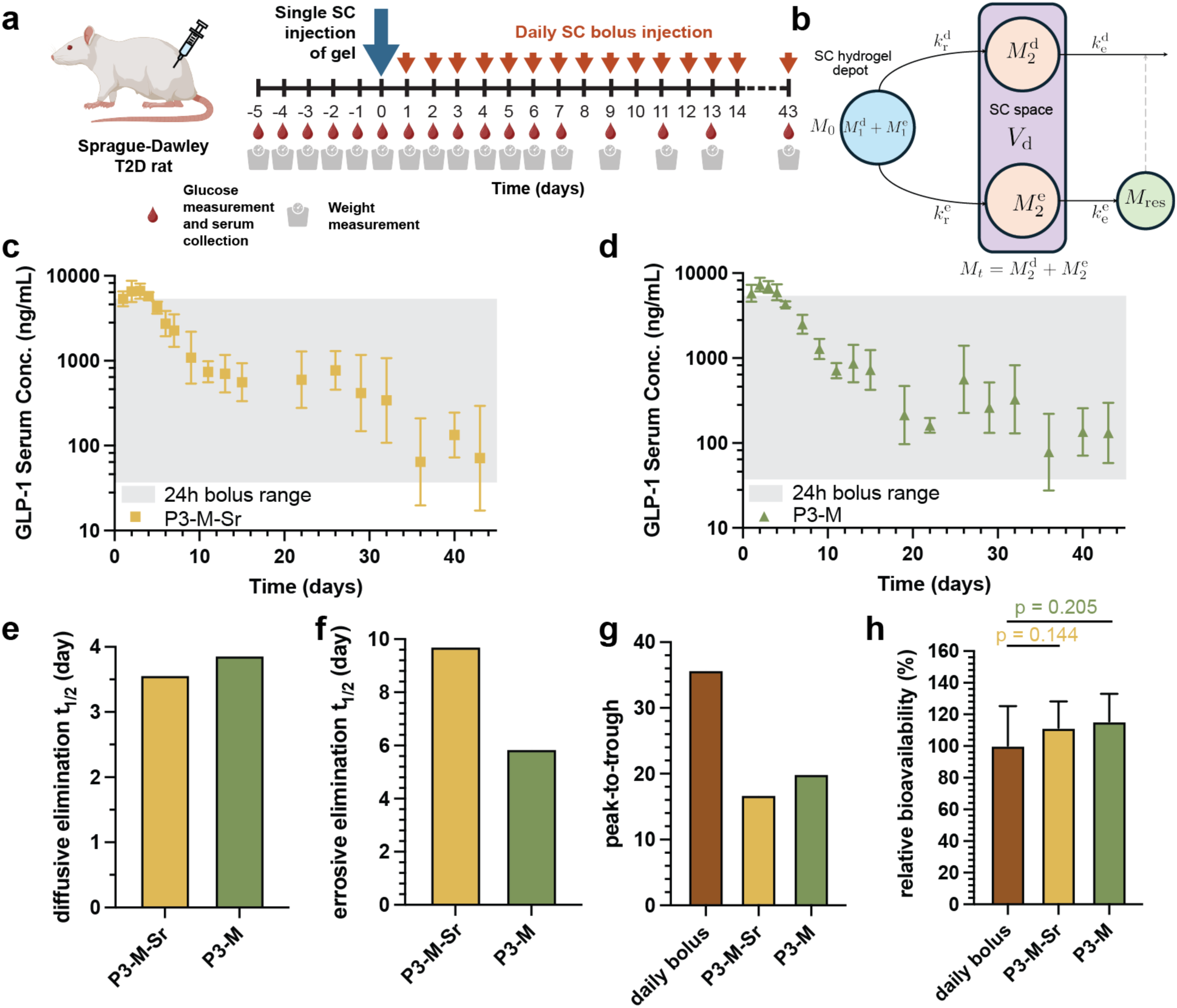
Pharmacokinetic evaluation of long-acting semaglutide depots in T2D rats over 42 days. **(a)** Study design for streptozotocin-nicotinamide-induced type 2 diabetic rats. A single subcutaneous hydrogel depot (1 mL, 900 μg) was administered on day 0. A control cohort received 42 daily injections of an aqueous Ozempic®-mimicking Sema formulation (20 μg daily; 840 μg total). Serum Sema, body weight, and glucose were monitored for 42 days. **(b)** Two-mode, two-compartment model used to extract diffusional and erosive elimination half-lives from serum concentration-time data. **(c–d)** Sema serum profiles for P3-M-Sr and P3-M formulations, with a gray box indicating the maximum and the minimum observed serum concentration from a 24-h semaglutide pharmacokinetics study in T2D rat.^21^ **(e–f)** Diffusional and erosive elimination half-lives derived from multi-modal two compartment model fitting (**Supplementary Discussion 4**). **(g)** Peak-to-trough ratios comparing burst versus sustained exposure. **(h)** Relative bioavailability values over the 42-day period benchmarked against the 20 μg daily bolus reference group. Methionine-containing depots provide prolonged exposure with reduced peak-to-trough fluctuation in the T2D model. Parameters shown in **(e–h)** were obtained from a combined fit across the full dataset. Statistical comparisons in **(h)** were performed using a two-tailed Welch’s t-test to account for unequal variances between groups. Methionine-containing depots provide prolonged exposure with reduced peak-to-trough fluctuation in the T2D model. Group size 5 ≤ n ≤ 7.

Based on our earlier burst-release assays, we selected P3-M-Sr and P3-M for evaluation in streptozotocin-nicotinamide-induced T2D rats. Methionine was prioritized because it consistently reduced burst release while extending sustained release, and its use as a sacrificial antioxidant in therapeutics is well established.^39,51^ The influence of Sr^2+^ on Sema bioavailability remained uncertain, warranting direct comparison. Rats received a single 1mL injection of depot-based Sema formulations (900 µg Sema), increased from 500 μL used in our previous studies.^21^ Serum Sema concentrations, body weight, and blood glucose were monitored over the 42-day treatment period. The glucose-lowering profile aligned with the pharmacokinetics, confirming that the depot-mediated release drives the observed pharmacodynamic response.

An initial evaluation of a P2 hydrogel formulation containing Tw20 (P2-0.6T20) yielded a pharmacokinetic profile characterized by a rapid early decline in serum Sema concentrations without a clearly defined C_max_ peak, reflecting a t_max_ shorter than 24 hours such that C_max_ occurred before the first sampling time point (**Supplementary Figure 3**). Moreover, this group exhibited poor bioavailability compared to our daily bolus control (**Supplementary Figure 3**). In contrast, our optimized formulations P3-M-Sr and P3-M (**Fig. 5c–d**) exhibited the characteristic shape of depot-based release, with an initial rise over the course of the first 3 days following administration, a relatively flat mid-phase (up to 3 weeks), and a multiweek decay resulting from slow erosion of the depot. These results demonstrated that increasing depot volume and network density shifts the release kinetics toward a delayed t_max_ and more sustained systemic exposure, enabling clearer resolution of C_max_ and improved pharmacokinetic control.

The C_max_ values observed for P3-M and P3-M-Sr formulations appear modest relative to bolus dosing, indicating that both depots effectively suppress burst release (**Fig. 5c–d**). P3-M-Sr shows a more gradual decline than P3-M, but both maintain measurable semaglutide levels through day 42. Model-derived half-lives of release and elimination (**Fig. 5e–f**) allow separating the contributions from passive diffusion-mediated versus depot erosion-driven loss. The two formulations produced similar diffusional half-lives (∼3.5 days), suggesting that early-phase transport is governed primarily by polymer-peptide interactions rather than Sr^2+^ modulation. In contrast, the erosive half-life differs noticeably between the groups, with P3-M-Sr erosively releasing cargo roughly 50% slower compared to P3-M, consistent with Sr^2+^ influencing late-stage retention by enhancing Sema multimerization and its interaction with the hydrogel matrix. Even so, both formulations exhibited erosion half-lives of exceeding 6 days. The gray shaded region in **Fig. 5c** and **5d** denotes the 24-hour pharmacokinetic window for each bolus administration of Sema in a T2D rat model, as previously reported.^21^ The upper and lower boundaries indicate the maximum and minimum serum concentrations measured within 24 hours of dosing, indicating that the P3-M and P3-M-Sr hydrogel-based formulations stay within the exposure range for standard Sema dosing for the duration of the study.

Analysis of the peak-to-trough ratio (**Fig. 5g**) provided a simple readout of exposure stability. The peak-to-trough ratios for both of our optimized depot formulations, P3-M-Sr (16.8) and P3-M (19.9), were more than two-fold lower than for the daily Sema bolus control (35.7) and more than seven-fold fold lower than our initial P2-0.6T20 formulation (142; **Supplementary Figure 1b**). This observation confirmed that the reduced burst release and gradual transition into sustained delivery for our optimized formulations is both highly desirable for tolerability and therapeutic efficacy.

A primary practical advantage of the methionine-containing hydrogel formulations is Sema stabilization, which can both blunt oxidation-driven early leakage and enable a smoother PK curve without sacrificing total exposure. Indeed, AUC measurements over the full 42-day period (**Fig. 5h**) indicated that both P3-M-Sr and P3-M formulations achieved excellent total systemic exposure with AUC values of (1.3±1.1)x10^6^ and (1.2±0.2)x10^6^ ng day mL-1, respectively. These values exceeded the total exposure for daily 20 µg bolus administrations over the treatment period ((1.0±0.4)x10^6^ ng day mL^-1^), commensurate with the higher total dosing for the study (*i.e.*, 900 µg total dose for hydrogel formulations and 840 µg total dose for the daily bolus control). The hydrogel dose was initially calibrated conservatively to offset anticipated exposure losses arising from incomplete release, local degradation, or injection-site retention. Normalization of the total hydrogel exposure by the total repeat-bolus control exposure indicates that these optimized hydrogel formulations enable comparable relative bioavailability to the bolus control (**Supplementary Figure 4**). Notably, the absence of a substantial reduction in AUC for the Sr^2+^-containing formulation indicates that concerns regarding compromised bioavailability were not supported *in vivo*.

These results demonstrated that hydrogel depot formulations comprising methionine maintain high total drug exposure for months following a single injection, with controlled early-phase behavior, low peak-to-trough fluctuation, and multi-week exposure in diabetic animals. Moreover, formulation with Sr^2+^ variant yields subtle differences in late-phase kinetics but no loss of total bioavailability. This long-term T2D rat study indicates that these hydrogel formulations represent a viable platform for stabilization and long-term release of lipidated incretin therapeutics in chronic disease models.

Following a general quarter-power scaling (**Supplementary Discussion 5**),^52^ the pharmacokinetics in humans can be estimated for analogous P3-M-Sr (**Supplementary Table 3**) and P3-M (**Supplementary Table 4**) formulations. For P3-M-Sr, we predict the half-lives to be: (i) diffusive release t_1/2_ = 2.77 days, (ii) diffusive elimination t_1/2_ = 13.15 days, (iii) erosive release t_1/2_ = 22.6 days, and (iv) erosive elimination t_1/2_ = 35.9 days. For P3-M, we predict the half-lives to be: (i) diffusive release t_1/2_ = 2.23 days, (ii) diffusive elimination t_1/2_ = 14.3 days, (iii) erosive release t_1/2_ = 50.9 days, and (iv) erosive elimination t_1/2_ = 21.7 days. These pharmacokinetic parameters can support dosing frequencies potentially reaching beyond 3-months in humans.^41,31^

### 2.5. Pharmacodynamic analysis in rats with type 2 diabetes

Pharmacodynamic evaluation following the treatment period of our study showed that hydrogel-based Sema formulations maintained relatively stable glucose levels throughout a 47-day period (**Fig. 6a**). Indeed, animals receiving the P3-M formulation remained within the normoglycemic range associated with normal postprandial fluctuations (80-180 mg dL^-1^), whereas daily bolus administration resulted in greater variability in glucose levels. Moreover, both hydrogel formulations significantly increased the percentage of study days during which animals remained within the normoglycemic range compared with daily bolus treatment (**Fig. 6b**). Another important marker, glycated hemoglobin (HbA1c), reflects average blood glucose levels over the preceding 2–3 months and is widely used as a clinical indicator of long-term glycemic control and risk of diabetes-related complications.^52^ Consistent with the observed blood glucose profiles, both hydrogel-based formulations exhibited HbA1c values after the treatment period comparable to daily bolus Sema dosing, with the P3-M formulation exhibiting less variable HbA1c values that trended lower than the control (4.4 ± 0.8 mmol mol^-1^ versus 6.1 ± 2.3 mmol mol^-1^; p=0.125). These results indicate that sustained Sema release from the hydrogel depot enables more stable glycemic control over extended periods relative to conventional bolus dosing.

**Fig. 6.**
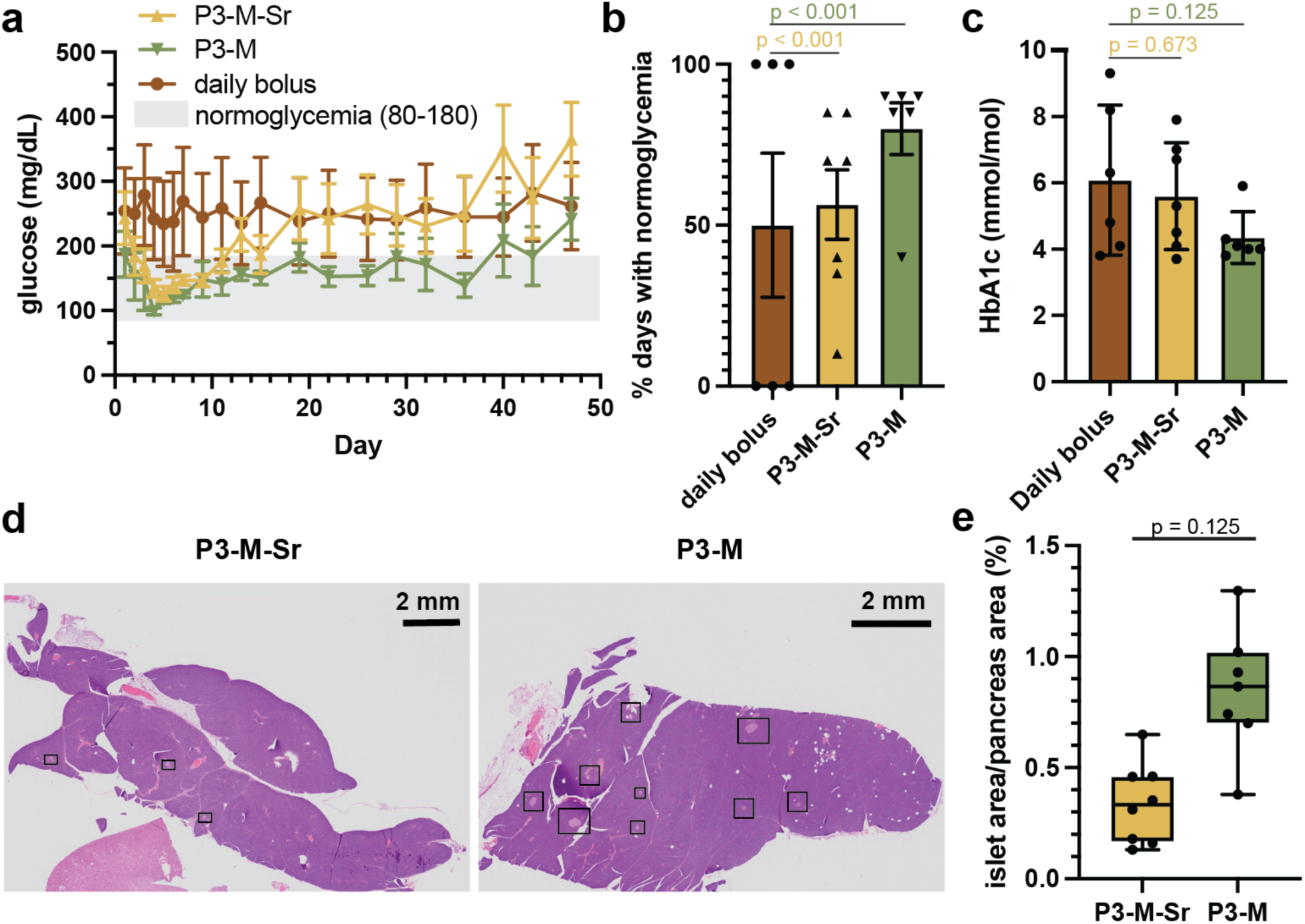
Pharmacodynamic outcomes and pancreatic remodeling following long-acting GLP-1 hydrogel treatments in T2D rats. **(a)** Blood glucose profiles over 6 weeks comparing daily bolus semaglutide, P3-M-Sr, and P3-M depots. The shaded region indicates the normoglycemic range (80–180 mg dL^-1^) **(b)** Percentage of study days during which animals remained within the normoglycemic range. **(c)** HbA1c levels measured at Day 47 following treatment. **(d)** Representative whole-pancreas H&E sections from untreated T2D, P3-M-Sr, and P3-M treated animals. Black boxes denote example regions used for islet quantification. **(e)** Quantification of islet-to-pancreas area ratios in P3-M-Sr and P3-M. Statistical significance was analyzed by Welch t-test for **(b), (c),** and **(e)** (5 ≤ n ≤ 7).

Normalized body weight change analysis further highlights differences in systemic exposure between treatment groups (**Supplementary Figure 5**). Animals receiving daily bolus Sema treatment exhibited a pronounced reduction in normalized body weight relative to expected vendor growth curves for healthy animals, with an average reduction of −35±2%, indicating substantial weight suppression during the study period. In contrast, rats treated with hydrogel formulations elicited normalized weight reductions of −19±3% for P3-M-Sr and −16±1% for P3-M reflecting more stable weight trajectories with significantly smaller deviations from expected growth. This distinction is important because the dosing strategy in this study was designed to maintain glycemic control in the T2D model rather than induce weight loss. The excessive weight reduction observed in the daily bolus group therefore reflects an undesirable pharmacodynamic effect associated with transiently high drug exposure. By contrast, the hydrogel formulations produced a more moderated physiological response while maintaining therapeutic efficacy, consistent with their sustained-release pharmacokinetic profiles. These results further support the advantage of hydrogel-based delivery for achieving stable metabolic control without excessive systemic exposure.

### 2.6. Tolerability and biodegradability of hydrogel depot formulations

Blood chemistry assessment on Day 0 pre-treatment and Day 47 post-treatment indicated overall metabolic and organ stability following treatment, suggesting excellent tolerability of these treatments (**Supplementary Figure 6**). Creatinine decreased in all treatment groups (**Supplementary Figure 6a**). For AST, an elevation was observed in the daily bolus control group, consistent with mild hepatic enzyme induction, while both hydrogel-based treatment groups showed no significant changes, indicating stable hepatic function (**Supplementary Figure 6b**). ALT remained stable across all treatments, with no evidence of hepatocellular injury (**Supplementary Figure 6c**). Bilirubin values were within normal limits in all groups, with P3-M-Sr showing a small but statistically significant decrease (p=0.0082, Welch’s t-test, 5 ≤ n ≤ 7) that remained well within the physiological range (**Supplementary Figure 6d**). For BUN, both P3-M-Sr and P3-M exhibited statistically significant decreases (p=0.018 and 0.0062, respectively), whereas the daily bolus control group showed consistent but nonsignificant levels, supporting good renal tolerance and absence of nephrotoxicity for all treatments (**Supplementary Figure 6e**). Although a subset of rats that received daily bolus dropped slightly below the physiological range, both hydrogel-based formulations showed modest, statistically significant reductions that remained within normal limits, potentially reflecting mild hydration effects rather than renal impairment.

To evaluate systemic and local biocompatibility of the sustained-release hydrogel formulations, the kidney, pancreas, liver, and skin at the injection site were examined 55 days post-administration (**Fig. 6d, Supplementary Figures 7–10, Supplementary Table 5**). In all treated animals, regenerative and occasionally degenerative lesions were observed within the pancreatic islets, characterized by mild to moderate cellular atypia (hypertrophy, increased basophilia, increased nuclear size, spindle-like morphology) and mild vacuolation of islet cells. Both hydrogel-treated groups exhibited mild regenerative islet changes consistent with adaptive remodeling in the diabetic state, without inflammatory cell infiltration. The P3-M treatment group exhibited a significantly higher islet area relative to total pancreas area compared to the P3-M-Sr treatment (p=0.0023, Welch’s t-test, 5 ≤ n ≤ 7), whereas the P3-M-Sr treatment group resembled untreated diabetic controls (**Fig. 6e, Supplementary Figure 9**). This increase in islet-organ-ratio suggests enhanced islet preservation or regeneration with P3-M treatment, likely driven by more consistent glucose control throughout the study period. These observations align with regenerative trends reported for GLP-1-based therapies,^53–55^ although specific endocrine cell populations cannot be resolved by routine histology.

At the subcutaneous injection site, depots persisted through the study with only mild inflammation and fibrosis, supporting good local tolerability and an intended multi-week release profile (**Supplementary Figure 10**). The P3-M depots were almost entire resorbed (one animal had no remaining hydrogel), which the P3-M-Sr depots exhibited more remaining material that was surrounded by a well-demarcated layer of foamy macrophages and low numbers of neutrophils and lymphocytes, accompanied by mild peripheral fibroblast proliferation and collagen deposition, which are features consistent with a controlled foreign-body response during gradual material resorption. No necrosis, abscess formation, or tissue destruction was observed in either hydrogel group, and the local tissue responses were comparable between P3-M-Sr and P3-M. As reported in other studies,^56^ the hydrogel was encapsulated by histiocytes and neutrophilic granulocytes with mild fibrosis, indicative of an ongoing but well-regulated degradative process. Consistent with our previous observations of HPMC-C_18_ in C57BL/6 mice, the present findings further support the ability of this hydrogel system to maintain structural integrity *in vivo* over extended durations (**Supplementary Figure 11**).^36^ The persistence of the depot demonstrated that residual Sema exposure detected at the end of the pharmacokinetic study is likely attributable to ongoing release, suggesting that these formulations can potentially support longer-term peptide release. Furthermore, these observations suggest that the injected hydrogel volume could potentially be reduced in future studies to specifically target monthly dosing.

Additionally, we conducted flow cytometry analysis of depots 4 days after treatment in C57BL/6 mice to quantify recruitment and infiltration of immune cells into the depots, including macrophages, neutrophils, and immune-cell derived reactive oxygen species (ROS) (**Supplementary Figure 12**). These studies demonstrated that no significant immune cell recruitment was observed across P3-noCargo, P3-Sema, P3-M-Sr, and P3-M formulations, suggesting that these hydrogel-based Sema formulations do not drive local inflammation or inflammatory cell recruitment. These observations corroborate our long-term studies in rats and indicate that these formulations are well tolerated.

## 3. Conclusions

Injectable depot technologies for the encapsulation and sustained release of lipidated peptide therapeutics are frequently constrained by low encapsulation efficiencies, burst release, mismatch in depot erosion and drug release, and erosional instabilities that create highly coupled formulation trade-offs driving extensive trial-and-error during development. In this work we report the development a dynamically crosslinked hydrogel depot technology enabling months-long exposure of lipidated peptides with complete bioavailability and lower exposure variability than a typical bolus injection. We also develop an engineering design framework to rationalize and reduce this complexity for this type of injectable hydrogel depot technology. Using Sema as a model lipidated peptide, formulation variables can be organized into three orthogonal control levers: (i) hydrogel vehicle formulation, (ii) cargo–excipient interaction, and (iii) oxidation stabilization, and each lever is linked to measurable structure–property–performance relationships across rheology, *in vitro* release, and *in vivo* pharmacokinetic and pharmacodynamic performance. This strategy transforms formulation from an open-ended optimization exercise into a guided decision process that narrows the experimental search space and accelerates identification of effective depot systems. Importantly, this design framework is not specific to semaglutide but applies broadly to formulation of amphiphilic or lipidated molecules in dynamic hydrogels.

Application of this framework demonstrated that lipidated peptides are intrinsically compatible with the C_18_-associated micellar crosslinks within our HPMC-C_18_ hydrogel network, enabling facile peptide encapsulation and sustained delivery without auxiliary surfactant excipients. While certain excipients can increase the size of the Sema constructs to improve retention within the hydrogel matrix, they may also destabilize the crosslinking within the hydrogel network and compromise pharmacokinetic control. Reversible, moderately labile multivalent coordination and pH modulation provide tunable routes to control peptide assembly in formulation to suppress early release while maintaining systemic availability. We also show that sacrificial antioxidant incorporation stabilizes Sema within the hydrogel formulations, mitigating early release and enabling smoother pharmacokinetics. Using a rat model of type 2 diabetes, we evaluated methionine-containing hydrogel depot formulations for sustained exposure after a single administration over the course of a 6-week study. These hydrogels exhibited a two-fold reduction in peak-to-trough fluctuation relative to daily bolus dosing. The improvement in consistency of exposure improves pharmacodynamic effects observed throughout the study, including glucose control, weight management, and pancreatic islet content that are maintained for over 47 days following treatment. Moreover, these hydrogel depot formulations are well tolerated according to blood chemistry and histopathology.

Overall, this work establishes an engineering-driven roadmap for depot formulation to reduce empirical iteration and enable a generalizable, predictive, mechanism-informed process for rational development of long-acting biologic formulations.

## 4. Materials and Methods

### Materials

#### Rheology

Rheological measurements were carried out on a TA Instruments DHR-2 stress-controlled rheometer equipped with a 20-mm serrated parallel plate geometry to mitigate wall slip artifacts. Samples were loaded at 600 µm geometry gap, and all tests were conducted at 21 °C. Oscillatory frequency sweep data were collected over a frequency range of 0.1–100 rad s⁻¹ using 1% strain amplitude, which falls within the linear viscoelastic regime. Strain-dependent behavior was assessed through amplitude sweeps at the frequency of 10 rad s⁻¹, spanning 0.01% to 10,000% of strain amplitude. Steady shear (flow) behavior was characterized by stress sweeps run from low to high stress under steady-state sensing conditions from shear rates of 0.1 to 100 s⁻¹. Yield stress was determined under stress-controlled conditions using a logarithmic progression of 10 points per decade.

#### In vitro assay for semaglutide sustained release in capillary tube

*In vitro* capillary tube release assay serves as an experimental model to evaluate hydrogel release behavior in an enzyme-free, quiescent saline environment (PBS). At each predetermined time point, all PBS buffer from the *in vitro* tube was collected and analyzed by ELISA to quantify the concentration of released drug in the collected PBS. New PBS is replenished when the sample from the previous timepoint is collected. Hydrogel samples (100 µL per formulation) were dispensed into four-inch-long glass capillary tubes. A 400 µL column of phosphate-buffered saline (PBS) was added above each sample to act as the release phase sink. At predetermined intervals (1, 3, 6, 12, 24, and 48 hours, as well as 1 and 2 weeks), the PBS layer was removed for analysis and replaced with fresh buffer to maintain sink conditions. Semaglutide concentrations in collected fractions were measured using a commercial ELISA kit (BMA Biomedicals, S-1530). Cumulative release curves were generated from triplicate samples (n = 3).

#### In vitro assay for semaglutide burst release

For each hydrogel formulation, 800 µL of PBS was added to 1.5 mL microcentrifuge tubes to serve as the incubation medium. Gel samples were delivered into the tubes using 21-gauge or comparable needle sizes, and 100 µL of each gel was dispensed to the bottom of the tube. Samples were incubated at 37 °C for 1.5 hours. Following incubation, the PBS supernatant was carefully aspirated and analyzed by semaglutide ELISA.

### Injection force measurements

Injection force measurements were conducted using a previously established protocol.^57,58^ A syringe pump was used to control the volumetric flow rate through a 1 mL BD syringe fitted with a 40 mm, 21-gauge needle. A calibrated load cell (FUTEK LLB300) mounted to the syringe pump recorded the axial force required for injection. Force–time data were collected using a LabVIEW acquisition program. Each measurement continued until the injection force reached a steady-state plateau, after which the pump was stopped.

### Animal studies protocol

Animal studies were performed with the approval of the Stanford Administrative Panel on Laboratory Animal Care (APLAC-32873) in accordance with NIH guidelines.

### Pharmacokinetics modeling for semaglutide PK in healthy mice

The *in vivo* pharmacokinetics of GLP-1 molecules released from subcutaneous gel administrations were quantified using a two-compartment model to obtain the release and elimination half-lives of the drug. The schematic representation of the model and equations are demonstrated in the supplementary information (**Supplementary Discussion 3**).

### Streptozotocin/Nicotinamide (STZ/NA) induced model of type-2 diabetes in rats

Male Sprague Dawley rats (8–10 weeks old, 230–350 g; Charles River) were used for T2D rat studies for evaluation of long-term pharmacokinetics and bioavailability of the GLP-1 RA hydrogel formulations under Stanford Administrative Panel on Laboratory Animal Care protocol #32873. Type 2 diabetes was induced using a combined nicotinamide (NA) and streptozotocin (STZ) regimen. Animals were fasted for 6–8 hours prior to induction. NA was prepared in sterile 1X PBS and administered intraperitoneally at 110 mg/kg. STZ was freshly dissolved in 10 mg/mL sodium citrate buffer and delivered intraperitoneally at 65 mg/kg. To reduce the risk of hypoglycemia following STZ dosing, rats were provided with a 10% sucrose solution in place of drinking water for the subsequent 24 hours. Blood glucose was monitored daily via tail vein sampling using a Contour Next handheld glucometer. Animals were classified as diabetic once they displayed at least three consecutive non-fasting glucose measurements between 140 and 290 mg/dL. The overall induction efficiency was approximately 50%.

### In vivo pharmacokinetics modeling for IV and SC in T2D rats

A 24-hour IV PK study was conducted to validate the PK parameters of Semaglutide in rats, and a single-compartment model was fit to obtain the elimination half-life (clearance rate) of the drugs. The differential equation governing the PK is:

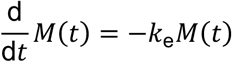

The IV PK data were fit to this equation. Here, *k_e_* is the elimination rate constant, from which the elimination half-life is determined as 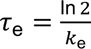. The SC gel data was also fit to the one-compartment model outlined above to obtain the half-lives of drug elimination from the body.

### In vivo pharmacokinetics and pharmacodynamics in T2D rats

For each treatment group (n = 6–8), baseline blood samples were collected from the tail vein on Day 0. Daily blood glucose levels were monitored using a handheld blood glucose monitor (Bayer Contour Next) for 42 days post-treatment.^59^ Blood glucose measurement was immediately followed by blood sampling from the tail vein for serum semaglutide quantification by competitive ELISA, performed daily during the first seven days and twice weekly thereafter. Plasma Sema concentrations were determined by ELISA at each time point, and total bioavailability was examined at the study endpoint. For the repeat bolus control group, sampling was conducted every 24 hours following each injection in the previous study.^21^

### Blood chemistry and histology analysis for biocompatibility

A blood chemistry panel was evaluated on Day 47 post injection. On Day 55, a total of sixteen T2D rats were euthanized using carbon dioxide and cardiac puncture for necropsy, including one T2D control rat without hydrogel injection, eight rats that received P3-M-Sr, and seven rats that received P3-m. The pancreas, liver, kidneys, and skin from the injection site were collected for histological evaluation. Tissues were fixed in 10% neutral-buffered formalin for 72 hours, trimmed and submitted to an external vendor for embedding and slide preparation (HistoTec Inc. Hayward, California). Tissue collection was performed at the Veterinary Service Center, Department of Comparative Medicine at Stanford University School of Medicine. Hematoxylin and eosin (H&E) staining was used to assess overall tissue morphology and local tissue response, while Masson’s trichrome staining was used to evaluate fibrosis at the injection sites. Histopathological examination was conducted by a veterinary pathologist (WR) blinded to treatment groups. For morphological examination, slides were examined with an Olympus BX53 microscope, Olympus DP27 Digital Camera and the respective cellSens Entry software, version 1.18 (Olympus America Inc. Waltham MA).

For the quantitative evaluation of the islet area, H&E-stained slides were digitized with the Olympus VS120 system and respective VS-ASW-S6 software, version 2.9.2 (Olympus America Inc. Waltham MA) at 40x magnification. The whole-slide images of pancreas were evaluated with the QuPath software, version 0.6.0.^60^ The whole pancreas area was detected by thresholding (Channel: Green, threshold: 170.0) and irrelevant tissues (e.g. lymph nodes) were excluded manually. Thereafter, islets were annotated manually. Single islet areas and whole organ area measurements were exported to Excel, summed up and islet-to-organ percentages were calculated. Data visualization and statistical analysis was performed using GraphPad Prism, version 10.6.1 (GraphPad Software LLC, Boston, MA).

### Peak-to-Trough analysis for T2D rat PK

Peak-to-trough calculations were performed by dividing the maximum observed plasma concentration (C_max_) from the long-term pharmacokinetic profile by the mean peptide concentration measured during the terminal steady-state period (days 11-42). Concentrations within this interval were averaged to obtain a representative trough value, as levels in this phase exhibit minimal day-to-day variation and reflect the sustained-release portion of the curve rather than early burst kinetics. This approach intends to provide a standardized metric to quantify depot stability, and the magnitude of concentration decline from peak to late-phase exposure.

### Pharmacokinetics modeling for T2D rat PK

The *in vivo* pharmacokinetics GLP-1 molecules released from subcutaneous gel administrations were quantified using a two-mode, two-compartment model to obtain the release and elimination half-lives of the drug, with the total release happening by a sum of diffusive and erosive mechanisms, and each mode having its characteristic release and elimination constants. A detailed schematic representation of this model and equations are demonstrated in the supplementary information (**Supplementary Discussion 4**).

### Quarter-power scaling for projected pharmacokinetics in humans from T2D rat PK

The cargo release and elimination half-lives obtained for Sema pharmacokinetics in T2D rats may be extrapolated to estimate the rates of clearance and persistence in humans using the “quarter-power” scaling law.^52^ Accordingly, the estimated respective half-lives in humans were calculated from the model fits to rat PK data, using t_human_ ∼ t_rat_(M_human_/M_rat_)^1/4^. The exact values are reported in the supplementary information (**Supplementary Tables 3 and 4**).

## 5. Data and materials availability

All data needed to evaluate the conclusions of this paper are present in the main text and/or the supplementary materials.

## 6. Code availability

All code supporting the findings of this study are available at: https://github.com/lyladong/pharmacokinetics-for-glp1-in-t2d-rats.

## 7. Acknowledgements

This work was supported by the National Institute of Diabetes and Digestive and Kidney Diseases (NIDDK) (R01DK119254). A.I.D. was supported by a Schmidt Science Fellows Award. C.D. is supported by James and Nancy Kelso Fellowship from the Stanford Interdisciplinary Graduate Fellowship program. L.T.N. received support from a Stanford Graduate Fellowship in Science and Engineering. In memoriam, we dedicate this work to L.T.N., whose unwavering commitment to advancing diabetes research, scientific rigor, and collaborative spirit left a lasting impact on this project and the broader scientific community. In addition, C.K.J., N.E., J.Y., O.M.S., C.M.W., and were supported by National Science Foundation Graduate Research Fellowships. S.K. gratefully acknowledges support from the 2023 Stanford SURF program and mentorship from Appel Lab. JHK is grateful for the support of the Agilent fellowship. We also thank all members of the Appel Lab for their valuable discussions and inputs throughout this project. The authors are grateful to the Stanford Animal Diagnostic Lab and the Veterinary Service Center staff for their technical assistance.

---

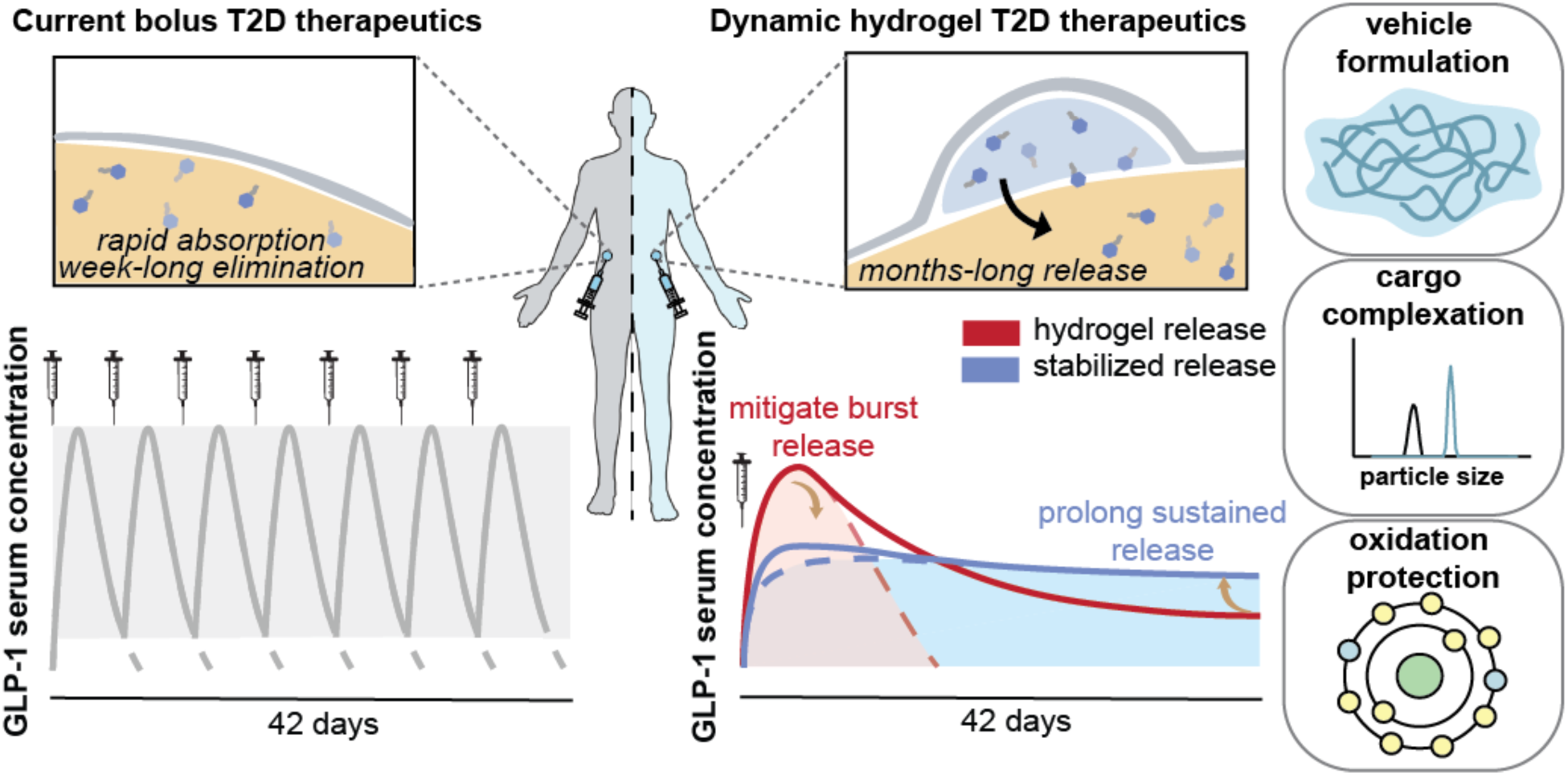
Schematic comparison of conventional soluble bolus injection and an injectable subcutaneous hydrogel depot for long-acting peptide delivery. Standard aqueous formulations of peptide therapeutics require frequent injections to maintain therapeutic exposure. Here, we develop an injectable supramolecular hydrogel depot technology that sustains systemic peptide levels for months following a single administration. By linking network mechanics, cargo association, and peptide stabilization, this work establishes a structure-property-performance framework for rational design of depot technologies over burst, diffusion, and erosion regimes to achieve long-acting peptide delivery.

