## Supplemental Information for "Structure–Property–Performance Engineering of Hydrogel Depots for Long-Acting Peptide Delivery"

**Supplementary Figure 7. Histological analysis of kidney samples. Kidney histology ... 22**
**Supplementary Figure 8. Histological analysis of liver samples. Liver histology ... 23**
**Supplementary Figure 9: Representative H&E–stained section of an entire pancreas from an untreated T2D rat ... 24**
**Supplementary Figure 10. Depot histology and study endpoint. ... 25**
**Supplementary Figure 11. C57BL/6 skin histology for P3 (HPMC-C18, no cargo) depot dissected after 10 days. ... 26**
**Supplementary Figure 12. Flow cytometry panel ... 27**

### Supplementary Discussion

#### Discussion S1. Continuous viscoelastic relaxation spectra

Continuous spectra of relaxation are used to analyze linear viscoelastic data, such as those obtained from frequency sweep measurements at small strains. For non-Maxwellian complex fluids, which includes most real fluids relevant in research, obtaining simpler rheological properties such as stiffness, plateau modulus, zero-shear viscosity, or a mean relaxation time is not straightforward. There exists a multi-modal, broad distribution of relaxation modes in real networks, and the analysis tool frequently employed is that of a spectral model description.[1, 2] A suitably chosen continuous spectrum model is fit to the data simultaneously for both *G*' and *G*", and the obtained spectra contain a sea of information about the relaxation processes in the material.

In this work, frequency sweep data obtained from SAOS tests were fit to continuous viscoelastic relaxation spectra models to obtain numerical values of rubbery plateau moduli and mean timescale of relaxation. In the continuous spectrum framework, viscoelastic moduli are defined as

$$G'(\omega) = \int_{-\infty}^{\infty} H(\tau) \frac{(\omega\tau)^2}{1 + (\omega\tau)^2} \mathrm{d} \ln \tau$$

$$G''(\omega) = \int_{-\infty}^{\infty} H(\tau) \frac{\omega\tau}{1 + (\omega\tau)^2} \mathrm{d} \ln \tau$$

where $H(\tau)$ is the continuous spectrum of relaxation, which can take analytical forms depending on the chosen distribution. From the viscoelastic "flow" component of the continuous spectra fits, the rubbery plateau modulus and the first mean relaxation time were calculated as

$$G_0 = \int_{-\infty}^{\infty} H_{\mathrm{fl}}(\tau) \mathrm{d} \ln \tau$$

$$\tau_{\mathrm{R}} = \frac{\int_{-\infty}^{\infty} \tau H_{\mathrm{fl}}(\tau)\mathrm{d}\ \ln\tau}{\int_{-\infty}^{\infty} H_{\mathrm{fl}}(\tau)\mathrm{d}\ \ln\tau}$$

where the flow component of the spectra, $H_{\mathrm{fl}}(\tau)$, is dependent on the choice of the relaxation spectrum chosen. For the materials tested in this study, the modified BSW spectrum was used,[3] which is given as

$$H(\tau) = H_{\mathrm{fl}}(\tau) + H_{\mathrm{gl}}(\tau)$$

$$H_{\mathrm{fl}}(\tau) = H_{\mathrm{f}}\left(\frac{\tau}{\tau_{\mathrm{max}}}\right)^{n_{\mathrm{f}}} \exp\left[-\left(\frac{\tau}{\tau_{\mathrm{max}}}\right)^{\beta}\right]$$

$$H_{\mathrm{gl}}(\tau) = H_{\mathrm{g}}\left(\frac{\tau}{\tau_{\mathrm{max}}}\right)^{-n_{\mathrm{g}}} \exp\left[-\left(\frac{\tau}{\tau_{\mathrm{max}}}\right)^{\beta}\right]$$

This spectrum has a stretched exponential cut-off at long times for both the flow and glassy terms, which works well for transient networks with dominant glassy contributions to stress relaxation and an incomplete terminal regime relaxation.

**Discussion S2. Diffusive release model for *in vitro* capillary release**

The cargo release data from *in vitro* capillary release assays were fit to the Ritger-Peppas model[4] to obtain power-law coefficients indicating the speed and nature of the diffusive release mechanism. The model is given as

$$\frac{M_t}{M_0} = kt^n$$

where $M_t$ is the cargo released over time $t$, $M_0$ is the initial cargo loading amount, $k$ is a release factor, and $n$ is the release coefficient.

**Discussion S3. Modeling semaglutide pharmacokinetics in C57BL/6**

The *in vivo* pharmacokinetics GLP-1 molecules released from subcutaneous gel administrations was quantified using a two-compartment model to obtain the release and elimination half-lives of the drug. The schematic representation of the model is shown below. The first compartment (I) is the subcutaneous (SC) hydrogel depot, which contains the protein cargo of total mass $M_0$. This compartment releases the cargo into the second compartment (II), the body (which includes the SC space, blood, etc.), and has a nominal volume $V_d$, also called the volume of distribution that can be assumed to be a volume of blood in a mouse. The kinetics of this release (I → II) are assumed to be first order, governed by the release rate constant $k_r$. The cargo is carried further from compartment I, metabolized, and eliminated from the body. The kinetics of this release (II → excretion) are again assumed to be of first order with an elimination rate constant $k_e$.

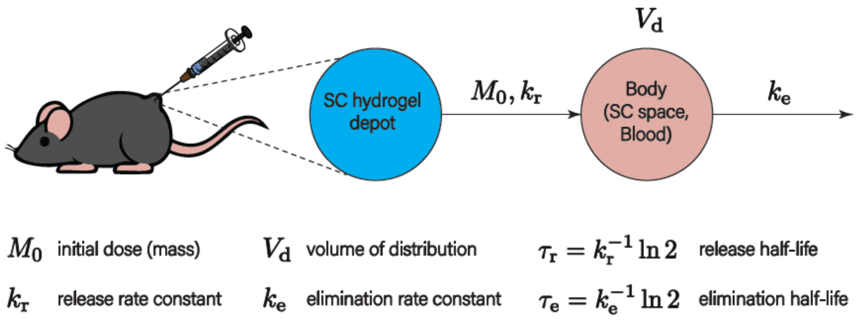

**Scheme S1: Two-compartment model for *in vivo* pharmacokinetics of semaglutide in mice.**

The mathematical form of this pharmacokinetics model consists of two kinetic balance equations of the mass of cargo for each compartment. The system of equations is given by

$$\frac{\mathrm{d}}{\mathrm{d}t} M_1 = -k_r M_1$$

$$\frac{\mathrm{d}}{\mathrm{d}t} M_2 = k_r M_1 - k_e M_2$$

where $M_1$ and $M_2$ are the mass of cargo in compartments I and II respectively. The two rate constants are obtained as fit parameters from fitting the model to PK data.

From the fits, the time at maximum concentration ($t_{max}$) and the maximum concentration ($c_{max}$) can be easily obtained. The AUC in the following pages is the area under the curve for the PK data.

**Discussion S4. Modeling pharmacokinetics in T2D rat model**

The *in vivo* pharmacokinetics GLP-1 molecules released from subcutaneous gel administrations was quantified using a two-mode, two-compartment model to obtain the release and elimination half-lives of the drug, with the total release happening by a sum of diffusive and erosive mechanisms, and each mode having its characteristic release and elimination constants. The schematic representation of the model is shown below.

The first compartment (I) is the subcutaneous (SC) hydrogel depot, which contains the protein cargo of total mass $M_0$. This compartment releases the cargo into the second compartment (II), the body (which includes the SC space, blood, etc.), and has a nominal volume $V_d$, also called the volume of distribution that can be assumed to be a volume of blood in a mouse. The kinetics of this release (I → II) are assumed to be first order, governed by the release rate constant $k_r$. The cargo is carried further from compartment I, metabolized, and eliminated from the body. The kinetics of this release (II → excretion) are again assumed to be of first order with an elimination rate constant $k_e$.

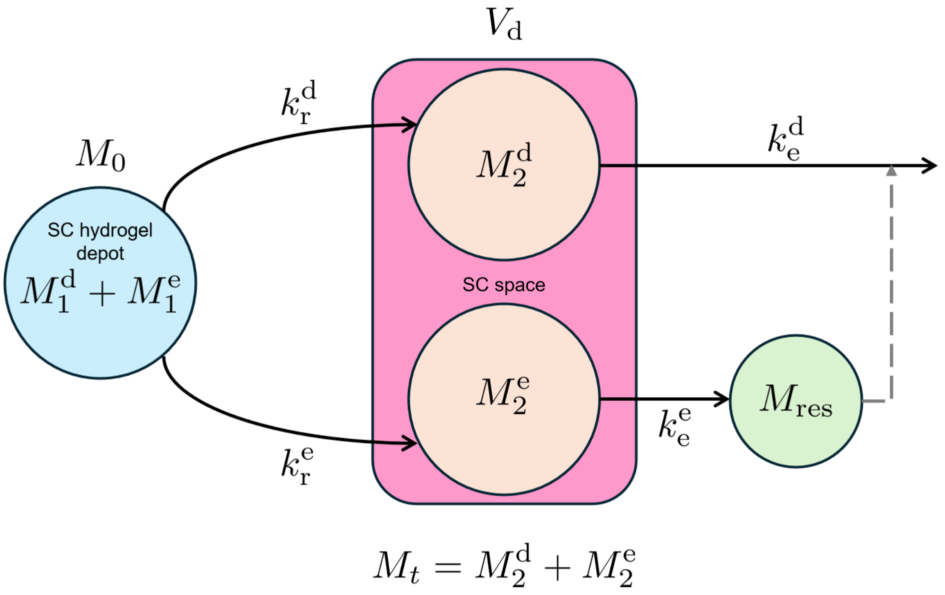

**Scheme S2: Two-mode, two-compartment model for *in vivo* pharmacokinetics of semaglutide in rats.**

The mathematical form of this pharmacokinetics model consists of two kinetic balance equations of the mass of cargo for each compartment, for both the diffusive and erosive modes. The system of equations for the diffusive mode is given by

$$\frac{\mathrm{d}}{\mathrm{d}t}M_1^{\mathrm{d}} = -k_{\mathrm{r}}^{\mathrm{d}}M_1^{\mathrm{d}}$$

$$\frac{\mathrm{d}}{\mathrm{d}t}M_2^{\mathrm{d}} = k_{\mathrm{r}}^{\mathrm{d}}M_1^{\mathrm{d}} - k_{\mathrm{e}}^{\mathrm{d}}M_2^{\mathrm{d}}$$

where $M_1^{\mathrm{d}}$ and $M_2^{\mathrm{d}}$ are the mass of cargo in compartments I and II respectively.

The system of equations for the erosive mode is given by

$$\frac{\mathrm{d}}{\mathrm{d}t}M_1^{\mathrm{e}} = -k_{\mathrm{r}}^{\mathrm{e}}M_1^{\mathrm{e}}$$

$$\frac{\mathrm{d}}{\mathrm{d}t}M_2^{\mathrm{e}} = k_{\mathrm{r}}^{\mathrm{e}}M_1^{\mathrm{e}} + k_{\mathrm{e}}^{\mathrm{e}}(M_{\mathrm{res}} - M_2^{\mathrm{e}})$$

where $M_1^{\mathrm{e}}$ and $M_2^{\mathrm{e}}$ are the mass of cargo in compartments I and II respectively; $M_{\mathrm{res}}$ is used to account for the residual (unreleased) cargo remaining in the hydrogel depot long after the diffusive process has reached steady state and only small vestiges of the depot persists at the injection site. The two rate constants for each mode are obtained as fit parameters from fitting the model to PK data. The half-lives are calculated as:

$$\tau_{\mathrm{r}}^{\mathrm{d}} = \frac{\ln 2}{k_{\mathrm{r}}^{\mathrm{d}}}$$

$$\tau_{\mathrm{e}}^{\mathrm{d}} = \frac{\ln 2}{k_{\mathrm{e}}^{\mathrm{d}}}$$

$$\tau_{\mathrm{r}}^{\mathrm{e}} = \frac{\ln 2}{k_{\mathrm{r}}^{\mathrm{e}}}$$

$$\tau_{\mathrm{e}}^{\mathrm{e}} = \frac{\ln 2}{k_{\mathrm{e}}^{\mathrm{e}}}$$

From the fits, the time at maximum concentration ($t_{\mathrm{max}}$) and the maximum concentration ($c_{\mathrm{max}}$) can be easily obtained. The AUC in the following pages is the area under the curve for the PK data.

**Discussion S5. Quarter-power scaling to project comparative human pharmacokinetics.**

To contextualize the long-acting pharmacokinetic behavior observed in rodents, allometric quarter-power scaling was applied to estimate projected pharmacokinetic parameters in humans [7]. Specifically, the half-lives of release and elimination may be extrapolated using the scaling law as

$$t_{1/2,\text{ human}} \approx t_{1/2,\text{ rat}} \left(\frac{M_{\text{human}}}{M_{\text{rat}}}\right)^{1/4}$$

where $M_{\text{human}}$ and $M_{\text{rat}}$ are the average masses of humans and rats respectively. For this study, we assumed the average mass of a human as 65 kg, for the average mass of a rat from the body weight data was taken to be 350 g. Accordingly, both release and elimination $t_{1/2}$ for diffusive and erosive modes obtained from fitting the rate PK data were scaled up.[5]

### Supplementary Tables

**Supplementary Table 1.** Naming scheme for formulations.

| Name | Sangelose (wt%) | Semaglutide (µg/mL) | PG (mg/mL) | Other excipients and notes | Buffer | pH | Injection volume (µL) |
|---|---|---|---|---|---|---|---|
| P3-M | 3 | 900 | 14 | methionine (1.5 mg/mL) | Tris-HCl 25Mm | 7.4 | 1000 |
| P3-M-Sr | 3 | 900 | 14 | methionine, SrCl2 (0.108 ug/mL) | Tris-HCl 25Mm | 7.4 | 1000 |
| P2-0.6T20 | 2 | 1800 | 14 | Tween20 | Tris-HCl 25Mm | 7.4 | 500 |
| Daily 20 µg | N/A | 40 | - | daily subcutaneous bolus, Rat | 1x PBS | 7.4 | 500 |
| P3 | 3 | 100 | 14 | - | 10 mM phosphate | 7.4 | 100 |
| P3@pH4 | 3 | 100 | 14 | - | 10 mM phosphate | 4 | 100 |
| P3-2T80 | 3 | 100 | 14 | Tween80 (20mg/mL) | 10 mM phosphate | 7.4 | 100 |
| P3-alb | 3 | 100 | 14 | rat serum albumin (1.5mg/mL) | 10 mM phosphate | 7.4 | 100 |
| P3-Zn | 3 | 100 | 14 | ZnCl2 (0.2mg/mL) | Tris-HCl 25Mm | 7.4 | 100 |
| P3-Zn@pH4 | 3 | 100 | 14 | ZnCl2 (0.2mg/mL) | Acetate buffer | 4 | 100 |
| P3-Sr | 3 | 100 | 14 | SrCl2 (0.12mg/mL) | Tris-HCl 25Mm | 7.4 | 100 |
| P3-Sr@pH4 | 3 | 50 | 14 | SrCl2 (0.12mg/mL) | Citrate buffer | 4 | 200 |
| P3-αT | 3 | 100 | 14 | α-tocopherol (0.5mg/mL) | Tris-HCl 25Mm | 7.4 | 100 |
| P3-αT-Sr | 3 | 100 | 14 | α-tocopherol (0.5mg/mL), SrCl2 (2.5mg/ml) | Tris-HCl 25Mm | 7.4 | 100 |
| P3-M | 3 | 100 | 14 | methionine (1.5mg/mL) | Tris-HCl 25Mm | 7.4 | 100 |
| P3-M-Sr | 3 | 100 | 14 | methionine (1.5mg/mL), SrCl2 (0.12mg/mL) | Tris-HCl 25Mm | 7.4 | 100 |
| P-CMC-Sr | 3 | 100 | 14 | Carboxymethyl Cellulose (25mg/mL), SrCl2 (0.022mg/mL) | Tris-HCl 25Mm | 7.4 | 100 |
| P-CMC-Ca | 3 | 100 | 14 | Carboxymethyl Cellulose (25mg/mL), CaCl2 (0.053mg/mL) | Tris-HCl 25Mm | 7.4 | 100 |
| Bolus 20 µg | - | 20 | - | single subcutaneous bolus, Mice | 1x PBS | 7.4 | 100 |

**Supplementary Table 2.** Pharmacokinetics parameters summary for formulations tested in healthy C57BL/6 mice.

| **Model:** | **C57BL/6, healthy female, 8-10 weeks** | | | | |
|---|---|---|---|---|---|
| **Name** | **release $t_{1/2}$ (day)** | **elim. $t_{1/2}$ (day)** | **AUC (ng d/mL)** | **$t_{max}$ (day)** | **$C_{max}$ (ng/mL)** |
| P3 | 0.4 | 1.27 | 22324 | 0.61 | 9241.3 |
| P3@pH4 | 1.05 | 1.17 | 29602 | 1.07 | 8735.1 |
| P3-2T80 | 0.3423 | 0.9319 | 22836 | 0.5734 | 12480 |
| P3-alb | 0.0584 | 0.4733 | 6297.2 | 0.1697 | 10795 |
| P3-Zn | 0.48 | 1.75 | 22096 | 0.86 | 8002.6 |
| P3-Zn@pH4 | 1 | 1.01 | 17694 | 0.99 | 6056.7 |
| P3-Sr | 0.4701 | 2.4849 | 39514 | 0.9971 | 11769 |
| P3-Sr@pH4 | 0.64 | 1.58 | 22754 | 0.92 | 7166.3 |
| P3-αT | 0.4946 | 2.3345 | 38370 | 0.9971 | 11585 |
| P3-αT-Sr | 0.4636 | 2.7546 | 43046 | 0.9971 | 12265 |
| P3-M | 0.3509 | 3.642 | 39540 | 0.9264 | 10668 |
| P3-M-Sr | 0.2657 | 4.129 | 40501 | 0.7852 | 10510 |
| P-CMC-Sr | 0.59 | 1.38 | 15673 | 0.92 | 5590.4 |
| P-CMC-Ca | 0.68 | 1.41 | 35668 | 0.92 | 11581 |
| Bolus 2 μg | 0.12 | 0.68 | 4777.8 | 0.31 | 3626.4 |

**Supplementary Table 3.** Pharmacokinetics parameters for P3-M-Sr, estimated as human pharmacokinetics followed by quarter scaling (Adjusted $R^2$ = 0.829).

| Fit parameters and results | | human (quarter scaling) |
|---|---|---|
| diffusive release $t_{1/2}$ (day) | 0.750 | 2.77 |
| diffusive elim. $t_{1/2}$ (day) | 3.56 | 13.2 |
| erosive release $t_{1/2}$ (day) | 6.11 | 22.6 |
| erosive elim. $t_{1/2}$ (day) | 9.72 | 35.9 |
| fraction of diffusive release | 0.871 | |
| residual cargo fraction (end) | 0.182 | |
| AUC | 52500 | |
| $t_{max}$ (day) | 1.55 | |
| $c_{max}$ (ng/mL) | 8550 | |
| AUC burst (7day)/total | 0.579 | |
| AUC diff/total | 0.642 | |
| peak/trough (at 42 day) | 16.8 | |

**Supplementary Table 4.** Pharmacokinetics parameters for P3-M estimated as human pharmacokinetics followed by quarter scaling (Adjusted $R^2$ = 0.848).

| Fit parameters and results | | human (quarter scaling) |
|---|---|---|
| diffusive release $t_{1/2}$ (day) | 0.622 | 2.30 |
| diffusive elim. $t_{1/2}$ (day) | 3.87 | 14.3 |
| erosive release $t_{1/2}$ (day) | 13.8 | 50.9 |
| erosive elim. $t_{1/2}$ (day) | 5.87 | 21.7 |
| fraction of diffusive release | 0.806 | |
| residual cargo fraction (end) | 0.0701 | |
| AUC | 51400 | |
| $t_{max}$ (day) | 1.55 | |
| $c_{max}$ (ng/mL) | 8920 | |
| AUC burst (7 day)/total | 0.617 | |
| AUC diff/total | 0.709 | |
| peak/trough (at 42 day) | 19.9 | |

**Supplementary Table 5.** Histology findings for individual reports.

| Rat ID | Group | Depot presence | Skin | pancreas |
|---|---|---|---|---|
| L0 | untreated T2D | N/A | - | - |
| L1 | P3-M-Sr | Yes | mild fibrotic | - |
| L2 | P3-M | Yes | mild fibrotic | - |
| L3 | P3-M-Sr | Yes | mild fibrotic | - |
| L4 | P3-M | Yes | mild fibrotic | - |
| L5 | P3-M-Sr | Yes | mild fibrotic | - |
| L6 | P3-M | Yes | mild fibrotic | - |
| L7 | P3-M-Sr | Yes | mild fibrotic | - |
| L8 | P3-M | Yes | mild fibrotic | Islet single cell necrosis |
| L9 | P3-M | Yes | mild fibrotic | - |
| L10 | P3-M-Sr | No | - | - |
| L11 | P3-M-Sr | Yes | mild fibrotic | - |
| L12 | P3-M | Yes | mild fibrotic | - |
| L13 | P3-M-Sr | Yes | mild fibrotic | mild interstitial fibrosis with mild infiltration of neutrophils |
| L14 | P3-M | Yes | mild fibrotic | mild focal nodular hyperplastic focus (incidental finding in aging rats) |
| L15 | P3-M-Sr | Yes | mild fibrotic | - |

### Supplementary Figures

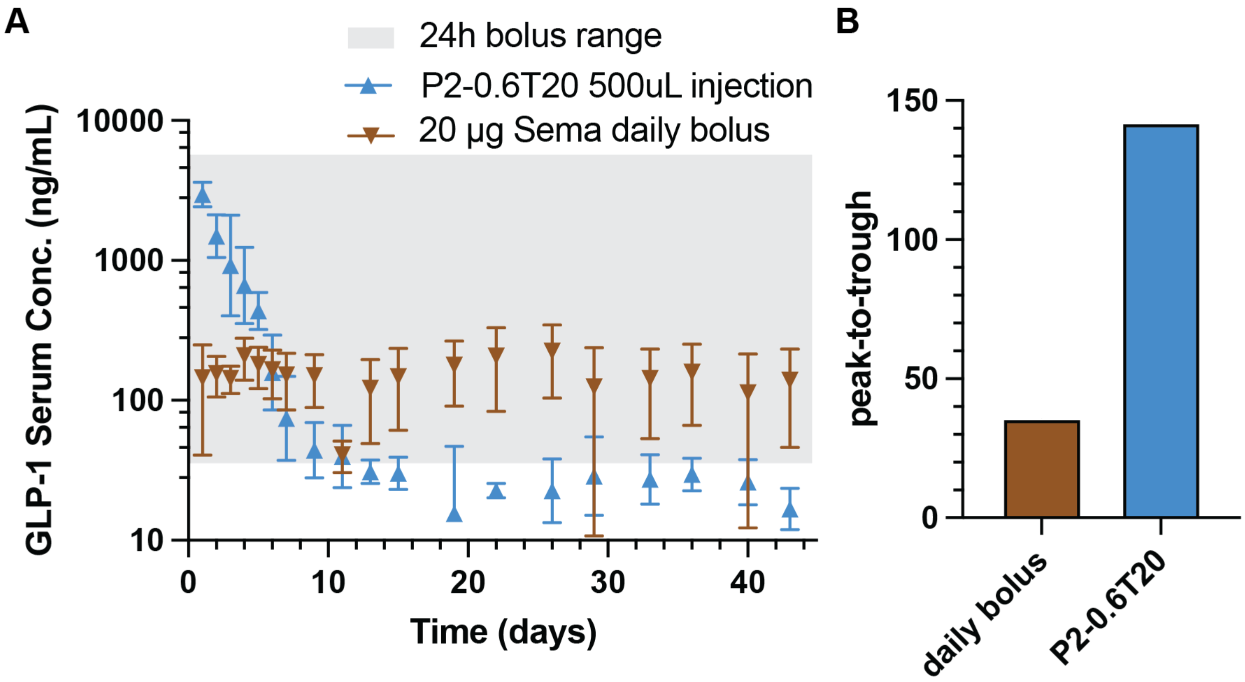

**Supplementary Figure 1. Pharmacokinetic profile of semaglutide in T2D rats following a single P2-0.6T20 hydrogel depot injection compared with daily bolus dosing.** (a) Serum GLP-1 concentrations over 42 days following a single 500 µL injection of P2-0.6T20 hydrogel depot (blue) compared to daily subcutaneous bolus dosing of semaglutide (20 µg, brown). The gray shaded region indicates the therapeutic concentration range achieved within 24 h after a single bolus injection. (b) Peak-to-trough ratio for daily bolus versus depot administration. Data for the daily bolus group and the 24 h bolus range are reproduced from a previously reported study.[6]

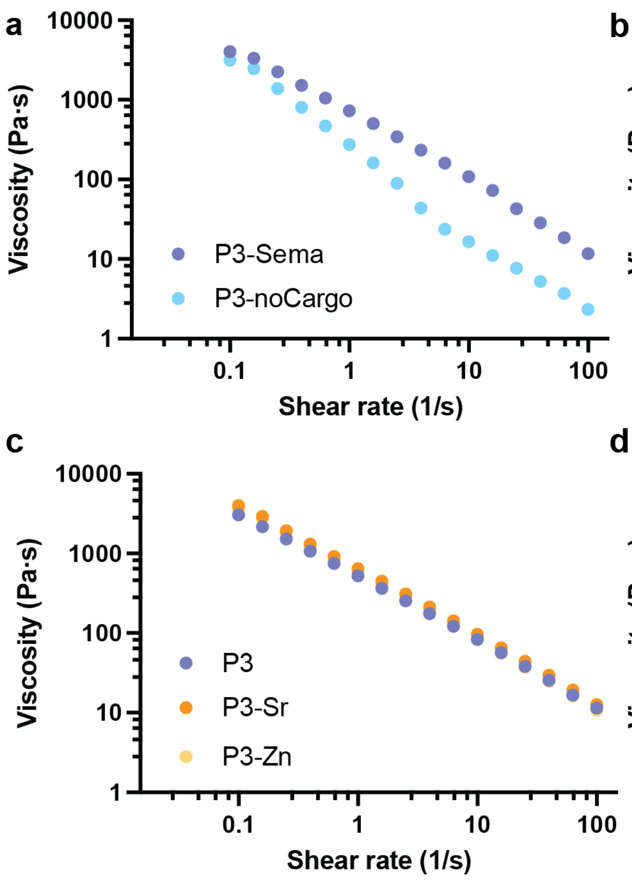

**Supplementary Figure 2. Flow sweep for depot formulations of interests.** Steady shear flow sweeps showing viscosity as a function of shear rate (0.1–100 $s^{-1}$) for the indicated formulations. **(a)** Comparison of P3-Sema and P3-noCargo demonstrates the effect of semaglutide incorporation on shear-thinning behavior. **(b)** Comparison of P3 and P3-alb evaluates the impact of albumin pre-complexation on network viscosity. **(c)** P3, P3-Sr, and P3-Zn highlight the influence of divalent cation complexation on flow response. **(d)** P3, P3-Sr, and P3-M-Sr compare combined complexation and antioxidant strategies. All formulations exhibit pronounced shear-thinning behavior, consistent with disruption of reversible hydrophobic junctions under increasing shear.

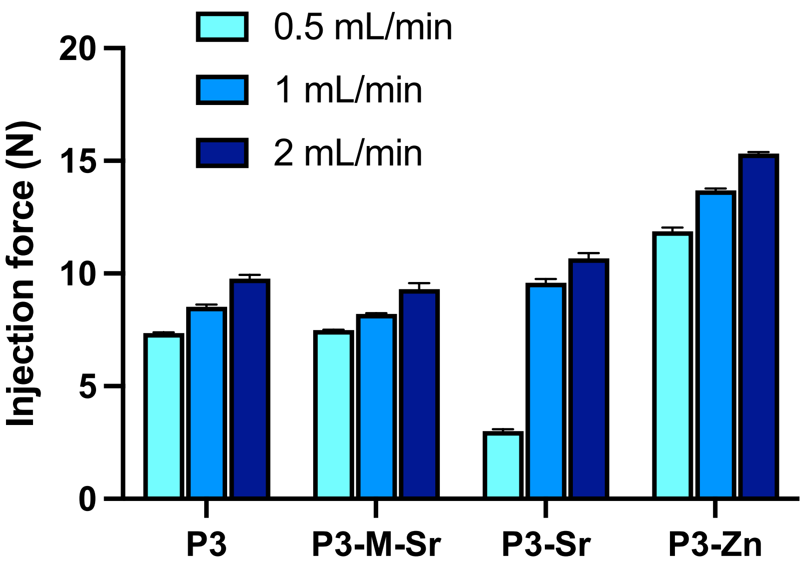

**Supplementary Figure 3. Injection forces of semaglutide depot formulations.** P3, P3-M-Sr, P3-Sr, and P3-Zn were measured at injection speeds of 0.5, 1, and 2 mL/min. All formulations exhibited forces within a reasonable range for subcutaneous administration. The addition of Zn markedly increased the injection force, suggesting stronger intermolecular or network interactions, whereas incorporation of Sr had minimal effect on injectability.

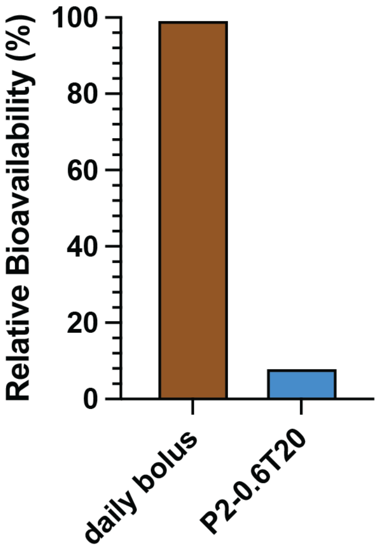

**Supplementary Figure 4. Relative bioavailability** normalized to daily bolus dosing. Data for the daily bolus group and the 24 h bolus range are reproduced from a previously reported study.[6]

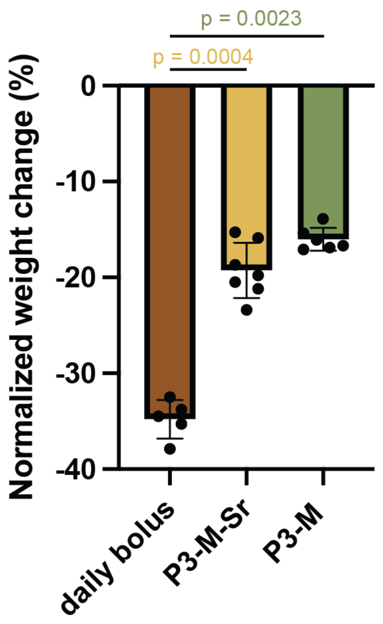

**Supplementary Figure 5. Normalized body weight change in T2D rats following treatment.** The control weights are from vendor's data for Sprague-Dawley rat standard growth data. Body weight change at Day 42 was calculated relative to baseline as fold change:

$$Fold\ change = \frac{Weight\ (Day\ 42)}{Weight\ (Day\ 0)}$$

To account for expected growth in untreated Sprague–Dawley rats, weight gain was normalized to the standard vendor growth curve using:

$$Normalized\ weight\ change\ (\%) = \frac{(Fold\ change\ treated) - (Fold\ change\ control)}{Fold\ change\ control}$$

where the control fold change represents expected growth based on vendor-reported Sprague–Dawley rat growth data. Data are shown for daily bolus semaglutide, P3-M-Sr hydrogel, and P3-M hydrogel groups. Points represent individual animals and bars indicate mean ± s.d. Statistical comparisons were performed against the daily bolus group using a two-sided Welch's t-test (p=0.0023 for P3-M-Sr and p=0.0004 for P3-M).

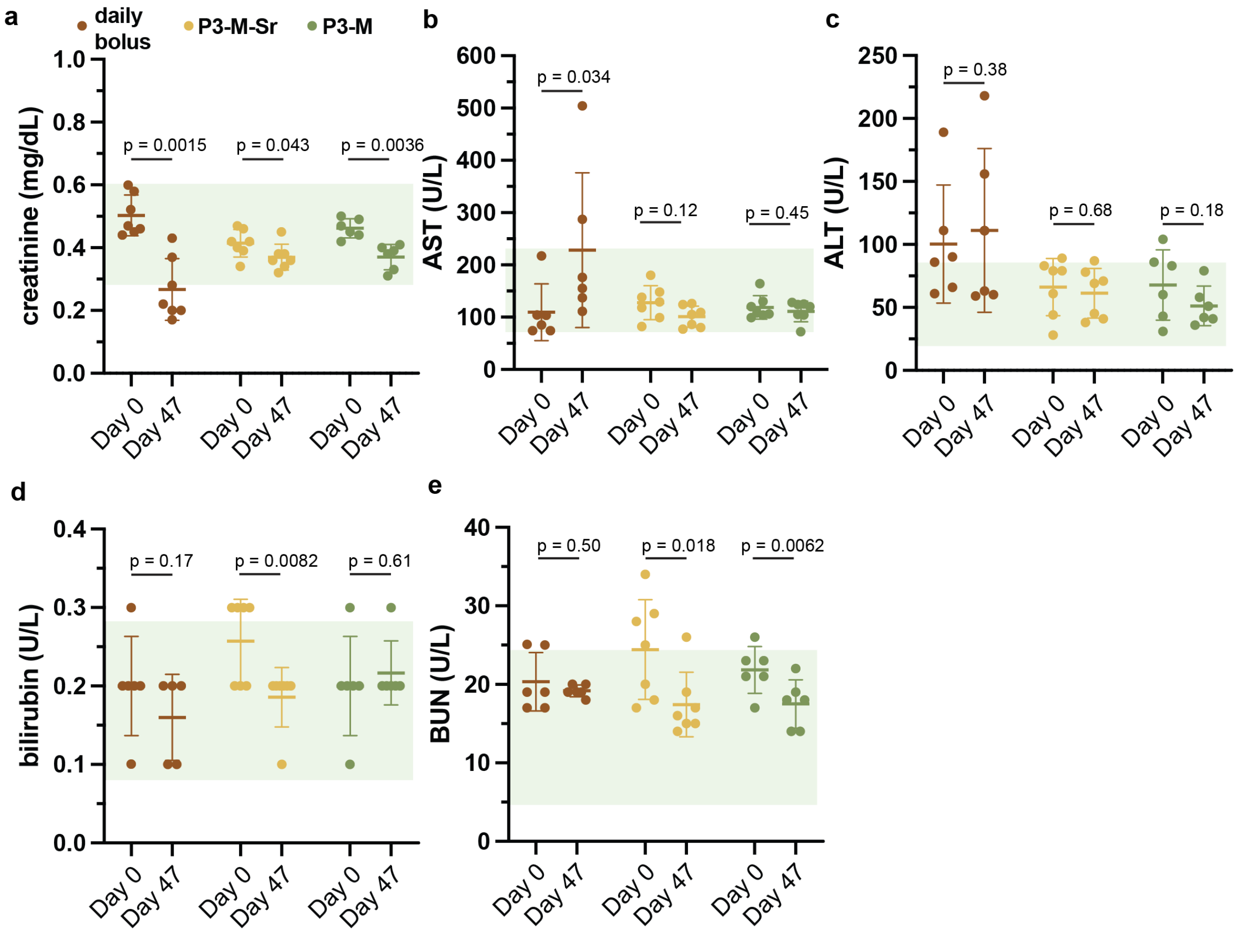

**Supplementary Figure 6. Blood chemistry panels** in streptozotocin–nicotinamide–induced T2D rats receiving daily bolus, P3-M-Sr, or P3-M hydrogel treatments, assessed at baseline (pre-treatment, Day 0) and at study endpoint (post-treatment, Day 47).

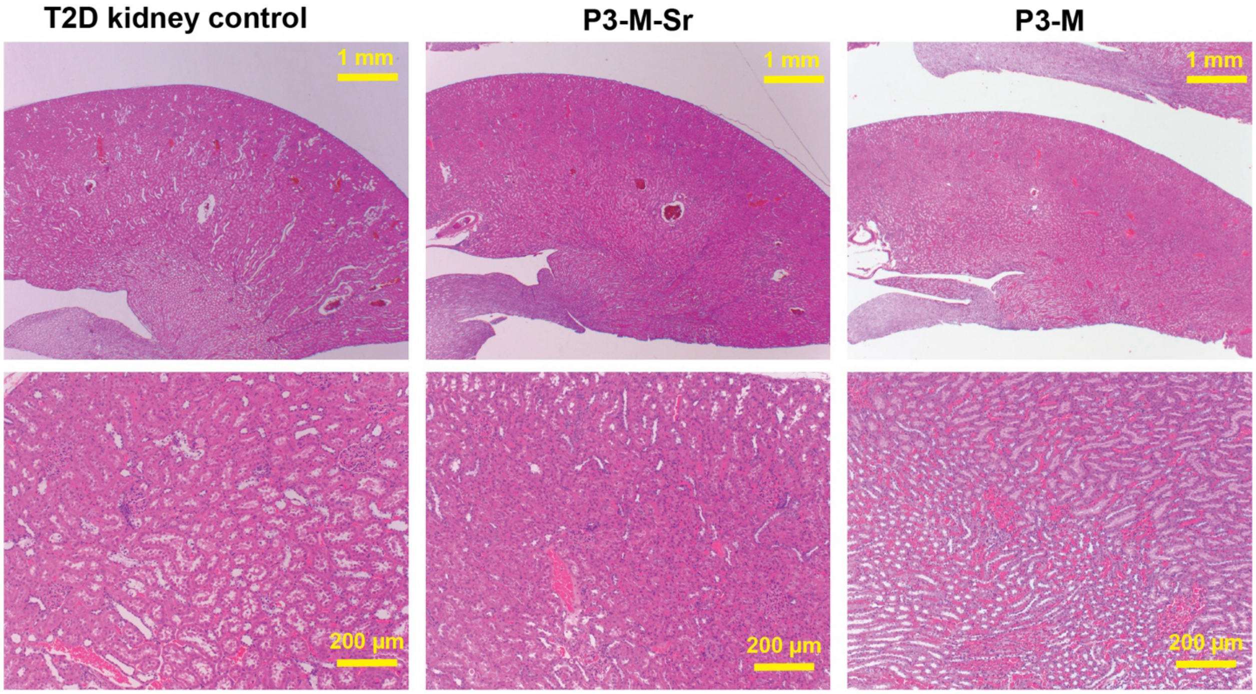

**Supplementary Figure 7. Histological analysis of kidney samples.** Kidney histology from streptozotocin–nicotinamide–induced T2D rats receiving daily bolus, P3-M-Sr, or P3-M hydrogel treatments, assessed at study endpoint (post-treatment, Day 55).

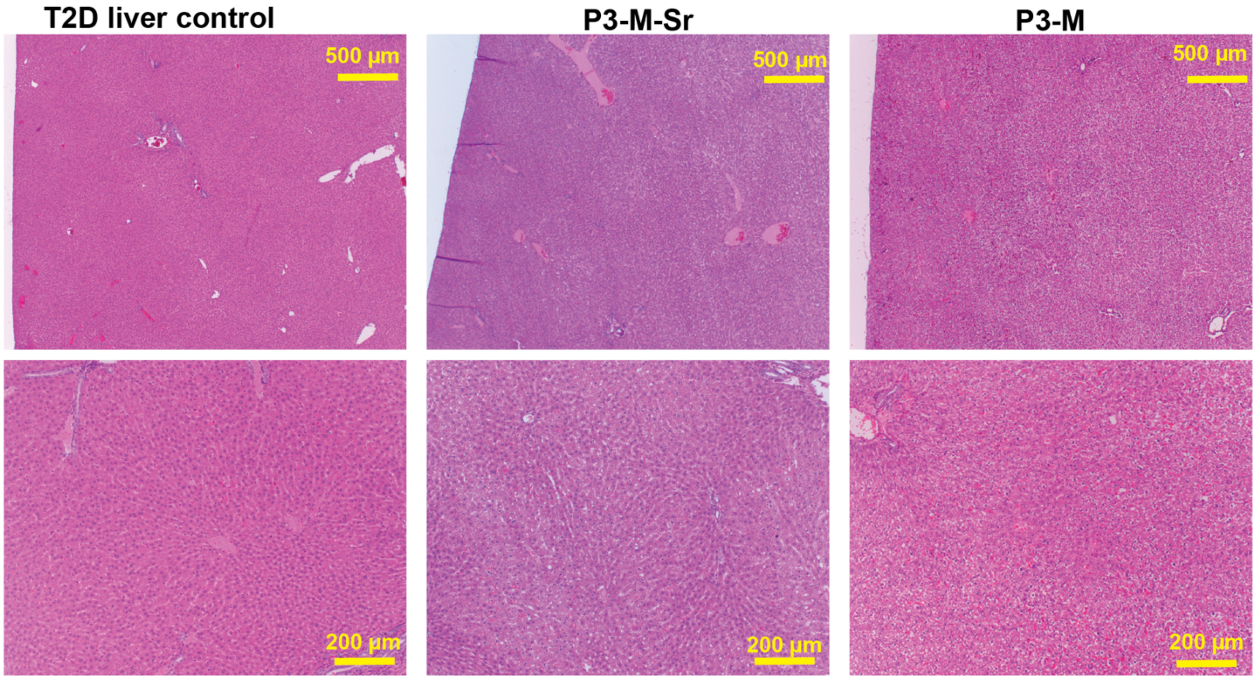

**Supplementary Figure 8. Histological analysis of liver samples.** Liver histology from streptozotocin–nicotinamide–induced T2D rats receiving daily bolus, P3-M-Sr, or P3-M hydrogel treatments, assessed at study endpoint (post-treatment, Day 55).

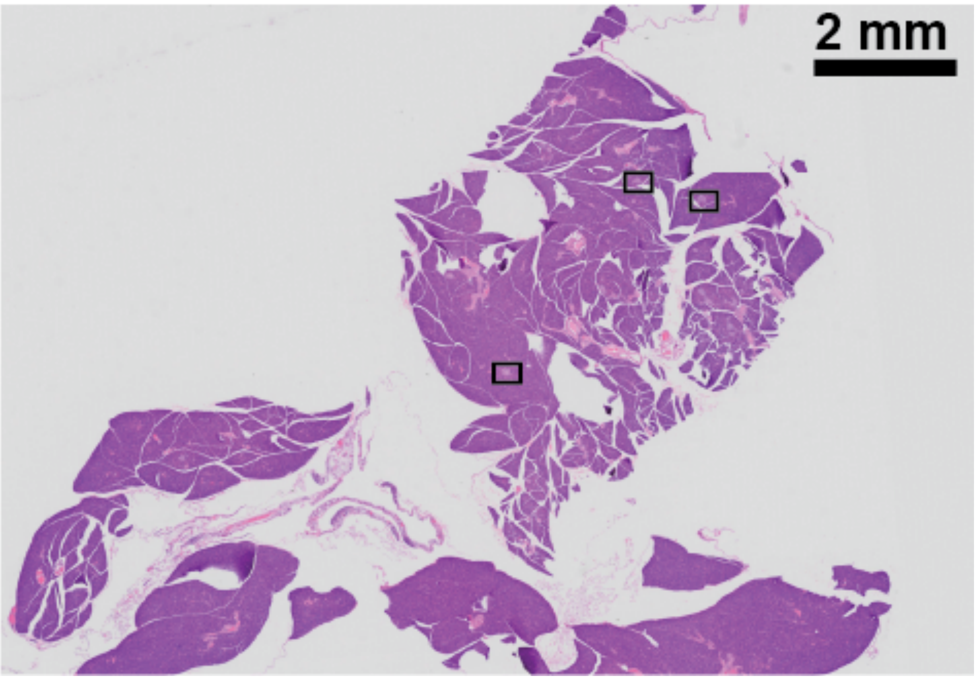

**Supplementary Figure 9: Representative H&E–stained section of an entire pancreas from an untreated T2D rat used for morphometric analysis of islet content.** Black boxes indicate representative regions containing pancreatic islets used for segmentation and area quantification. The islet area was determined relative to the total pancreatic tissue area, yielding an islet area fraction of 0.368% (islet area / pancreas area) for this specimen.

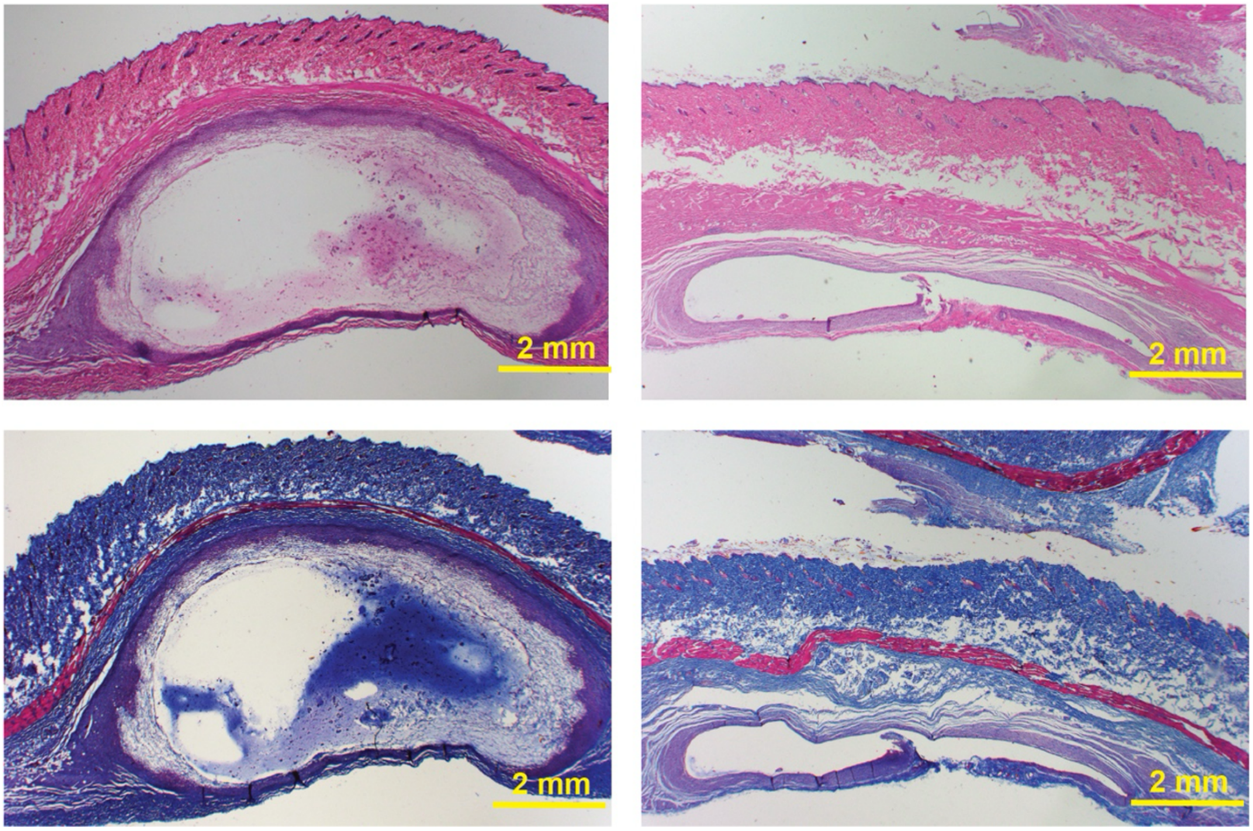

**Supplementary Figure 10. Depot histology and study endpoint.** Representative histological images of the P3-M-Sr (left) and P3-M hydrogel depots taken at the endpoint of the study and stained with either hematoxylin & eosin (H&E; top) or Masson's Trichrome (bottom). These images indicate that the P3-M hydrogel depot is almost completely eroded away while some of the P3-M-Sr hydrogel depot persists. No adverse tissue reactions or fibrosis were observed in either depot group.[7]

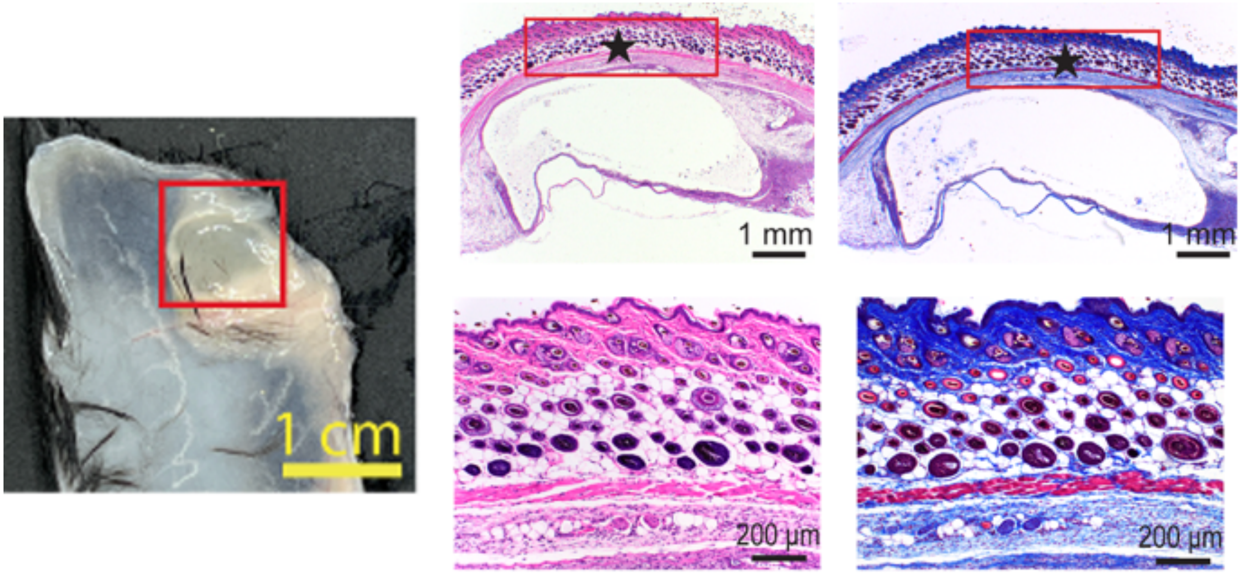

**Supplementary Figure 11. C57BL/6 skin histology for P3 (HPMC-C18, no cargo) depot dissected after 10 days.** (left top and bottom: hematoxylin & eosin (H&E); right top and bottom: Masson's Trichrome). Healthy female C57BL/6 were injected 100 $\mu$L of 3wt% HPMC-$C_{18}$ and gel was harvested at Day 10 for histological analysis that showed the presence of subcutaneous depot.[7] Representative photograph of the P3 depot (left). Histological images of the P3 depot stained with either hematoxylin & eosin (H&E; middle) or Masson's Trichrome (right) indicate there is no adverse tissue reactions or fibrosis.[7]

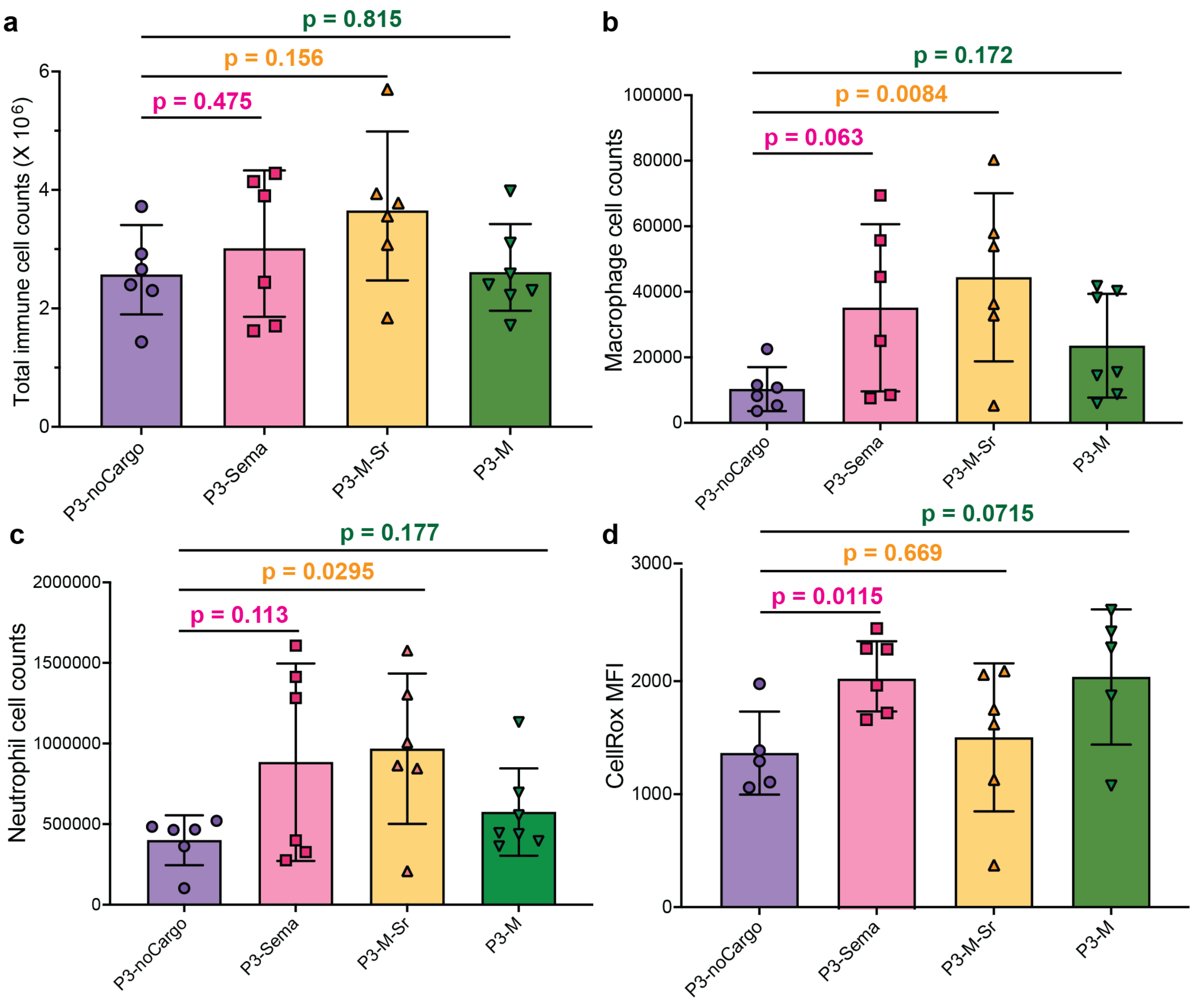

**Supplementary Figure 12. Flow cytometry panel for depot harvested from C57BL/6 at day 4 post injection,** including (a) total immune cell counts, (b) macrophage cell counts, (c) neutrophil cell counts, and (d) immune cells reactive oxygen species (ROS) mean florescence intensity (MFI) (Welch's t-test).
